# Dendritic, delayed, and stochastic CaMKII activation underlies behavioral time scale plasticity in CA1 synapses

**DOI:** 10.1101/2023.08.01.549180

**Authors:** Anant Jain, Yoshihisa Nakahata, Tetsuya Watabe, Polina Rusina, Kelly South, Kengo Adachi, Long Yan, Noriko Simorowski, Hiro Furukawa, Ryohei Yasuda

## Abstract

Behavioral time scale plasticity (BTSP), is a form of non-Hebbian plasticity induced by integrating pre- and postsynaptic components separated by behavioral time scale (seconds). BTSP in the hippocampal CA1 neurons underlies place cell formation. However, the molecular mechanisms underlying this behavioral time scale (eligibility trace) and synapse specificity are unknown. CaMKII can be activated in a synapse-specific manner and remain active for a few seconds, making it a compelling candidate for the eligibility trace during BTSP. Here, we show that BTSP can be induced in a single dendritic spine using 2-photon glutamate uncaging paired with postsynaptic current injection temporally separated by behavioral time scale. Using an improved CaMKII sensor, we saw no detectable CaMKII activation during this BTSP induction. Instead, we observed a dendritic, delayed, and stochastic CaMKII activation (DDSC) associated with Ca^2+^ influx and plateau 20-40 s after BTSP induction. DDSC requires both pre-and postsynaptic activity, suggesting that CaMKII can integrate these two signals. Also, optogenetically blocking CaMKII 30 s after the BTSP protocol inhibited synaptic potentiation, indicating that DDSC is an essential mechanism of BTSP. IP3-dependent intracellular Ca^2+^ release facilitates both DDSC and BTSP. Thus, our study suggests that the non-synapse specific CaMKII activation provides an instructive signal with an extensive time window over tens of seconds during BTSP.

## Introduction

Synaptic plasticity is the basis of acquiring and storing new information in the brain ^1^. Synaptic plasticity can be induced by specific patterns of electrical activity such as high-frequency synaptic stimulation ^2,3^, spike-timing-dependent plasticity (STDP) ^4–6^, or synaptic stimulation coupled with various neuromodulators ^7,8^. Ca^2+^ influx via the postsynaptic activation of NMDA receptors (NMDARs) and the downstream synapse-specific activation of CaMKII is the central mechanism that leads to an increase in the dendritic spine volume and AMPAR number and conductance, causing synaptic potentiation ^9–13^. In the classical Hebbian mechanism, coordinated pre- and postsynaptic activation leads to a relieve of the Mg^2+^ block in NMDAR, allowing Ca^2+^ ions to flow into dendritic spines.

Despite the extensive cellular and molecular understanding of synaptic plasticity, particularly at hippocampal CA3-CA1 synapses, there is limited direct evidence of specific plasticity mechanisms that occur at these synapses during behavioral learning ^14^. Computational studies suggest that the Hebbian plasticity rules that applies to the milliseconds require modification to explain simple learning behaviors that occur over seconds to minutes ^15^. Recently, a behavioral time scale plasticity (BTSP) paradigm was discovered at CA3-CA1 synapses and was suggested to be involved in CA1 place cell formation ^16,17^. In this plasticity, brief postsynaptic depolarization is paired with presynaptic inputs within a “behavioral time scale” or hundreds of milliseconds to seconds to induce synaptic potentiation during place cell induction. Synaptic potentiation can be induced either by forward pairing (presynaptic stimulation followed by depolarization) or converse pairing (depolarization followed by presynaptic stimulation). BTSP can be interpreted as a product of 1) eligibility trace, an input-specific signal that lasts for a few seconds, and 2) instructive signals by postsynaptic depolarization, which induces synaptic potentiation to the eligible synapse. For the converse BTSP (cBTSP), the instructive signal also needs to influence the synapse-specific signal over a few seconds. Studies suggest that the BTSP requires plateau potential and Ca^2+^ spikes in dendrites, perhaps providing instructive signals ^17,18^. However, the molecular representations of the eligibility trace and the instructive signal are unknown.

Using 2-photon fluorescence lifetime imaging (2pFLIM) of fluorescent resonance energy transfer (FRET) sensor, previous studies have shown that Ca^2+^ Calmodulin-dependent kinase II (CaMKII), a kinase critical for long-term synaptic plasticity ^11^, is activated in a synapse-specific manner during glutamate uncaging-induced structural plasticity of dendritic spines ^19,20^. CaMKII remains active for 1-6 seconds after glutamate uncaging due to autophosphorylation at the T286 site ^20,21^. This time scale of CaMKII activation (seconds) makes it a potential entity for the eligibility trace.

In the current study, we investigated the role of CaMKII in BTSP using whole-cell electrophysiology, glutamate uncaging, and 2pFLIM imaging of an improved CaMKII conformational sensor in hippocampal slices. First, we developed a glutamate uncaging protocol to induce BTSP at individual dendritic spines. Second, to image CaMKII activity with high sensitivity, we improved the sensitivity of a CaMKII sensor by ∼2 fold. Using this sensor, we did not find any evidence that CaMKII is activated during the BTSP protocol. Instead, a dendritic, delayed, and stochastic CaMKII activation (DDSC) occurs several tens of seconds after the induction of BTSP. We confirmed the requirement of DDSC by inhibiting CaMKII using optogenetic CaMKII inhibitor paAIP2 ^22^ at different time points after BTSP induction. Finally, we found that both DDSC and BTSP require IP3-dependent Ca^2+^ release from internal stores. Our experiments demonstrate the critical role of non-synapse-specific CaMKII activation as an instructive signal spanning an extended time scale (tens of seconds) in BTSP. Furthermore, BTSP appears to be induced by integrating two different time scales, one for integrating pre- and postsynaptic inputs over behavioral time scale to induce DDSC (seconds) and the other for integrating input-specific priming signal with DDSC over tens of seconds.

## Results

### Behavioral time scale plasticity can be induced in single spines at proximal apical dendrites

Previous studies have shown that during place cell formation, activity localized at CA1 dendrites precedes the somatic cell firing suggesting synapse-specific activation of plasticity ^23^. Thus we investigated whether behavioral time scale plasticity (BTSP), which is one of the mechanisms described for place cell formation ^17^, can be induced in single spines. To do so, we employed a whole-cell patch-clamp electrophysiology on CA1 neurons in organotypic hippocampal slices and measured 2-photon uncaging-evoked excitatory postsynaptic potentials (EPSPs) from 1-2 spines of secondary branches of proximal apical dendrites before and after the induction of BTSP (**Fig. 1a**). To induce BTSP, we delivered a train of 5 uncaging pulses on one spine at 1 Hz intervals, and then after a 750 ms delay from the last pulse (3.25 s from the center of uncaging pulses), gave a 600 pA current injection pulse for 300 ms (**Fig. 1b**). We found that this protocol induced a 93±16% (n=19) potentiation in EPSP amplitude in the stimulated spines (**Fig. 1d, e**), but not in the adjacent spines (13±10%, n=11, **Fig 1d, e**). We also developed a converse BTSP protocol (cBTSP), in which a current injection (600 pA for 300 ms) was delivered 750 ms before 5 uncaging pulses at 1 Hz (**Fig 1c**). This protocol also induced a similar EPSP potentiation (81±25%, n = 15) in EPSP amplitude in stimulated spines but not in adjacent spines (13±10%, n = 12) (**Fig. 1f, g**). Similarly, in acute hippocampal slices, the BTSP protocol also induced potentiation (103±29%, n = 12) in the stimulated spines but not in adjacent spines (18±7%, n = 8) (**Fig. 1h-j**). These results demonstrate that BTSP can be induced in single dendritic spines in a synapse-specific manner both in acute and organotypic slices.

**Figure 1:**
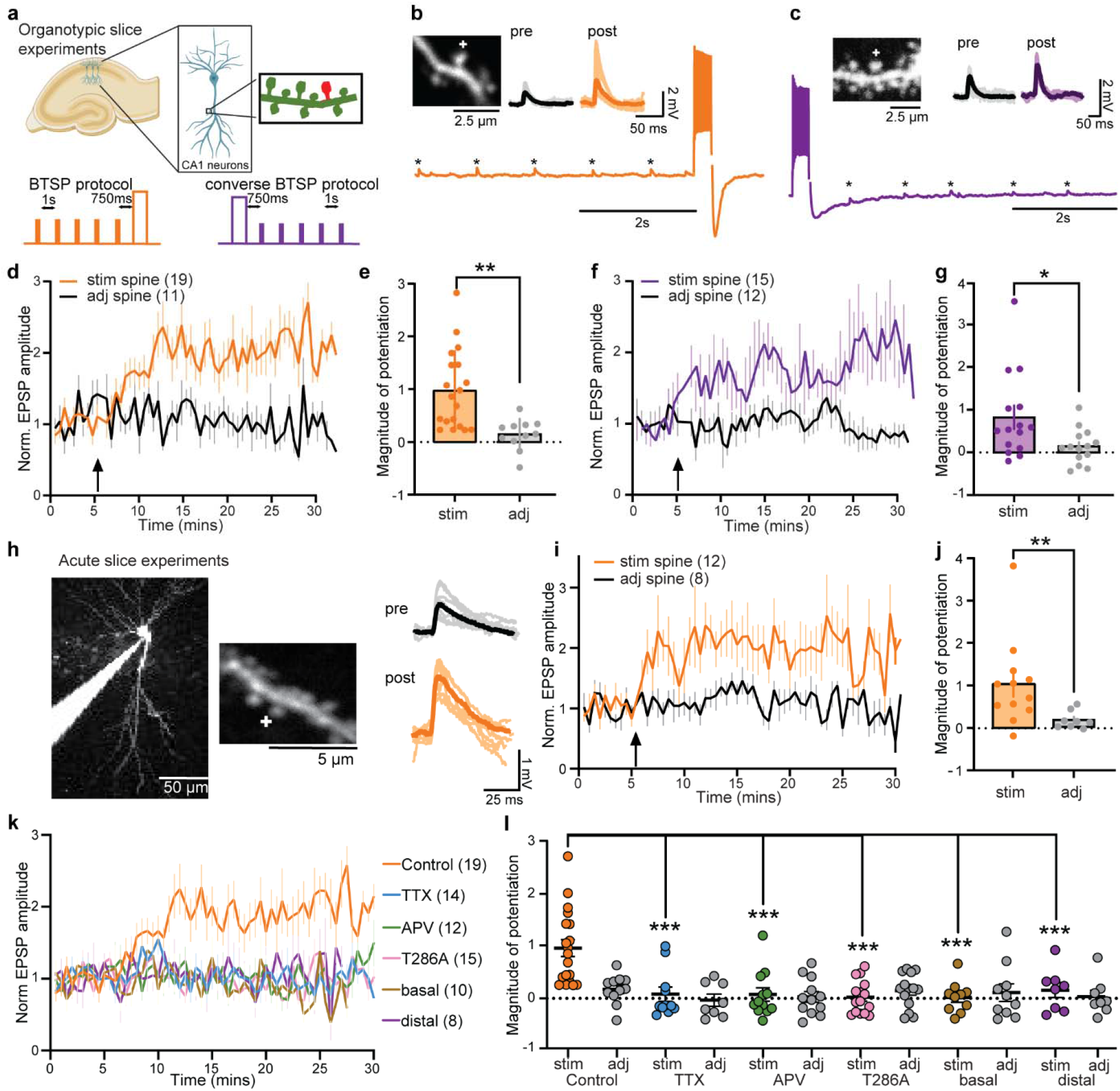
Glutamate uncaging protocols to induce BTSP at proximal apical CA1 dendritic spines. **a,** Top: Schematics of the experimental setup. Bottom: BTSP and converse BTSP (cBTSP) protocols: five uncaging pulses were given at an individual spine before (orange) or after (purple) a 600 pA current injection (300 ms) with a delay of 750 ms. **b,** A representative image of a dendrite of a CA1 neuron in an organotypic hippocampal slice (right top), uncaging-evoked EPSPs (10 recording and average) before and after BTSP from the spines on the dendrite (left top), and electrical trace during the BTSP protocol (bottom). **c,** Same as b, but with the cBTSP protocol. **d-e,** Mean time course (**d**) and summary of the magnitude of potentiation (25-30 min) (**e**) of uncaging-evoked EPSP amplitude normalized to the baseline (0-5 min) at stimulated spines (n=19, orange) and adjacent spines (n=11, black) in response to BTSP protocol (arrow). **p < 0.01, two-tailed t-test. **f-g**, Same as c-d, but with cBTSP protocol (n = 15 for stimulated spines and 12 for adjacent spines). *p < 0.05, two-tailed t-test. **h,** Left: Representative Alexa 594 filled image of a CA1 neuron and a dendritic shaft in acute hippocampal slices, where BTSP was induced. Right: Uncaging evoked EPSP traces of the stimulated spine before (10 EPSPs) and after (10 EPSPs) induction of BTSP. **i,** Averaged time course of normalized EPSP amplitude in response to BTSP induction in stimulated and adjacent spines in acute hippocampal slices. Numbers of cells are in the figure. **j,** Summary of the magnitude of potentiation of EPSP amplitude. *p < 0.05, two-tailed t-test, p<0.05. **k-l,** Average time course (**k**) and the summary (**i**) of the magnitude of BTSP-induced EPSP potentiation under various conditions. TTX and APV are bath applied. T286A is data from *Camk2a*^T286A^ mice. Basal and distal: BTSP protocol on basal or distal (> 200 μm from the soma) dendrites, respectively. Sample numbers (spines) are in the figure. Two-way ANOVA with multiple comparison test (Dunnett’s correction) ***p<0.001.

Since basal and distal dendrites receive inputs different from proximal apical dendrites ^24^, we investigated whether BTSP can also be induced in these dendrites. Notably, the same BTSP protocol failed to induce EPSP potentiation at basal or distal apical dendrites (<200 μm from the soma) (basal: 7±17% n = 10, distal, 12±12%, n = 8) (**Fig. 1k, l**, **Extended Fig. 1**). Thus, BTSP induction mechanism may depend on the location of dendritic branches. Next, we examined the molecular mechanism of BTSP using pharmacological inhibitors and transgenic mice. We found that voltage-gated sodium channel inhibitor TTX (1 μM) and NMDAR inhibitor APV (50 μM) inhibit BTSP induction in stimulated spines (**Fig. 1k, l**, **Extended Fig. 1**). Furthermore, mutant mice in which CaMKII activity is reduced (αCaMKII^T286A^) ^25^ showed no BTSP (**Fig. 1k, l**, **Extended Fig. 1**). These results suggest that, similarly to the Hebbian LTP, postsynaptic spiking and the activation of NMDAR and CaMKII are required for BTSP induction.

### Improved CaMKII sensor detected delayed global CaMKII activity after BTSP induction

If CaMKII represents the eligibility trace, its activation should be localized in the stimulated spines and last for a few seconds. This synaptic input will not release the Mg^2+^ block; thus, Ca^2+^ signaling is likely to be small. To detect anticipated small CaMKII activity during BTSP, we optimized our CaMKII sensor by putting two accepters in the original Green Camuiα sensor (2dV-Camui)^19^ (**Fig. 2a, Supplementary Note**). We found that 2dV-Camui displays a 2-fold higher signal with a similar time course compared to the previous version when tested in cell lines (**Fig. 2b-d**, **Extended Fig. 2-1**), and uncaging-evoked CaMKII activation in dendritic spines (note that structural LTP experiments in Fig. 2 are under zero extracellular Mg^2+^ condition. All other experiments are performed with 1 mM Mg^2+^) (**Fig. 2e-g**) ^19^. As expected, we observed a fast decay in CaMKII activation for a sensor in which the critical autophosphorylation site (T286) is mutated ^21^ (**Fig. 2h**, **i**). Also, mutations that turn off calmodulin binding (T305 and T306) eliminated its activation (**Fig. 2h**, **Extended Fig. 2-2**). Moreover, we confirmed that this sensor forms the usual dodecameric holoenzyme based on fluctuation correlation spectroscopy (FCS) and fluorescence-coupled size-exclusion chromatography (FSEC) (**Extended Fig. 2-3**, **2-4**).

**Figure 2:**
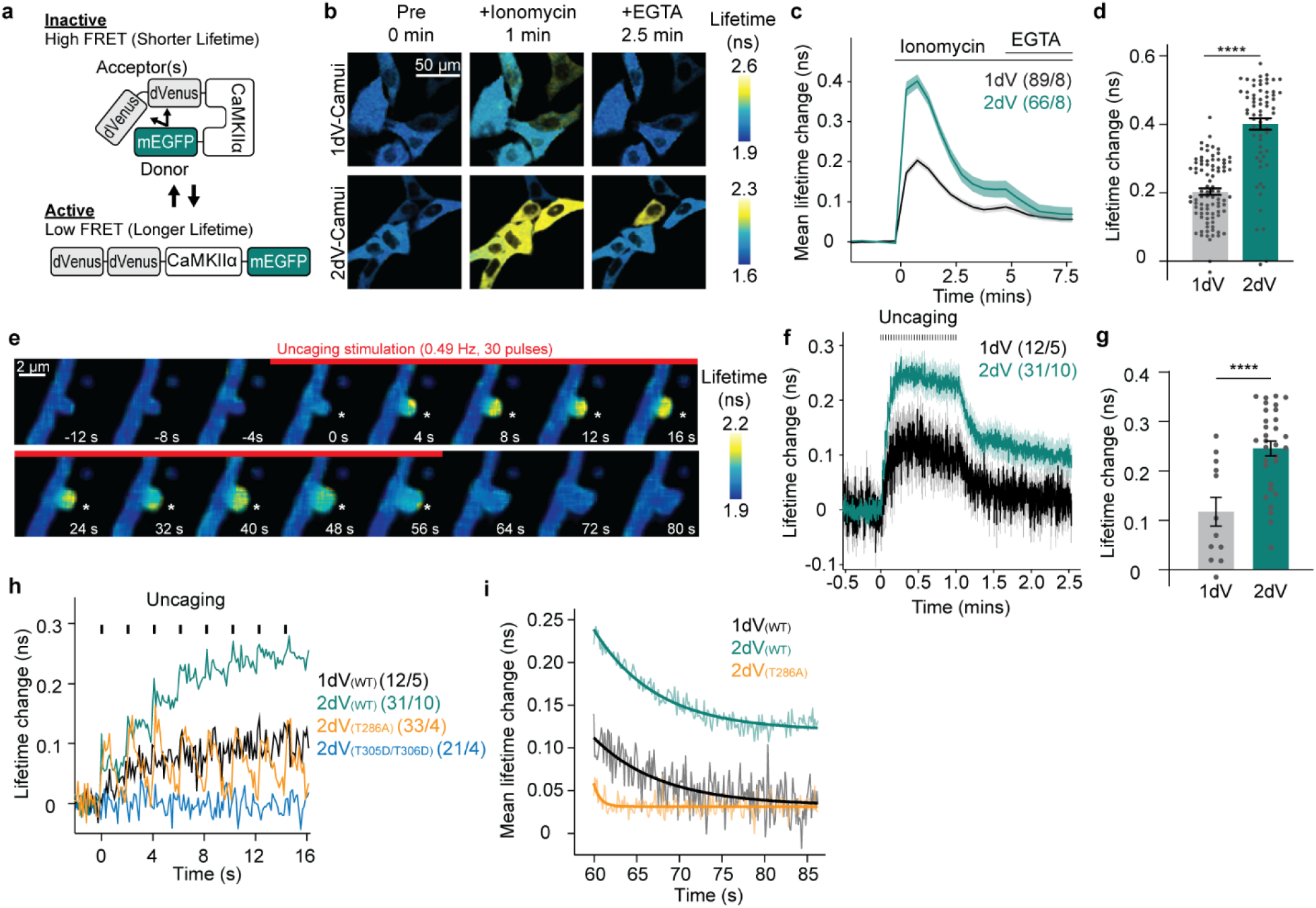
Optimization and characterization of novel conformational CaMKII FRET sensor. **a,** A schematic of Camuiα-2dv CaMKII sensor where N- and C-termini of CaMKIIα are labeled with donor EGFP and 2 dimVenus acceptor fluorophores. Activation of CaMKII results in a conformational change and a decrease of FRET efficiency. **b,** Fluorescence lifetime images of HeLa cells expressing Camuiα-2dV (top) or original Green Camui with 1 dimVenus (1dV-Camui) (bottom) before, during ionomycin and during EGTA. **c-d,** Averaged time course (**c**) and summary of the peak (1 min post drug application) (**d**) of fluorescent lifetime changes of 2dV-Camui (2dV) and Green Camui (1dV) in response to bath application of ionomycin (3 µM) in HeLa cells. 2dV-Camui showed ∼2 fold higher sensitivity. Numbers of samples are in the figure (cells / cultures). **e,** Fluorescence lifetime images of CA1 dendrites in hippocampal culture before, during, and after glutamate uncaging at 0.49 Hz in zero extracellular Mg^2+^. **f-g,** Averaged time course (**f**) and summary of the peak (6-11^th^ uncaging pulses) (**g**) of fluorescence lifetime changes in stimulated spines and adjacent dendritic shafts. The numbers of samples are in the figure (spines / cells). **h,** Closer view of the lifetime change during first 8 uncaging pulses in 2dV-Camui wildtype (2dV WT), Green-Camui wildtype (1dV WT), 2dV-Camui T286A, and 2dV-Camui T305D/T306D. The numbers of samples are on the figure (spines/cells). **i,** The decay kinetics in spines after uncaging. The fitting curve indicate single exponential fitting (y = Aexp(-t/τ) + B, where the fast time constant (τ) is 7.9, 7.3 and 0.74 s for 1dV WT, 2dV WT, and 2dV T286A, respectively. *** p < 0.001, **** p < 0.001, Two-tailed t-test.

**Figure 3.**
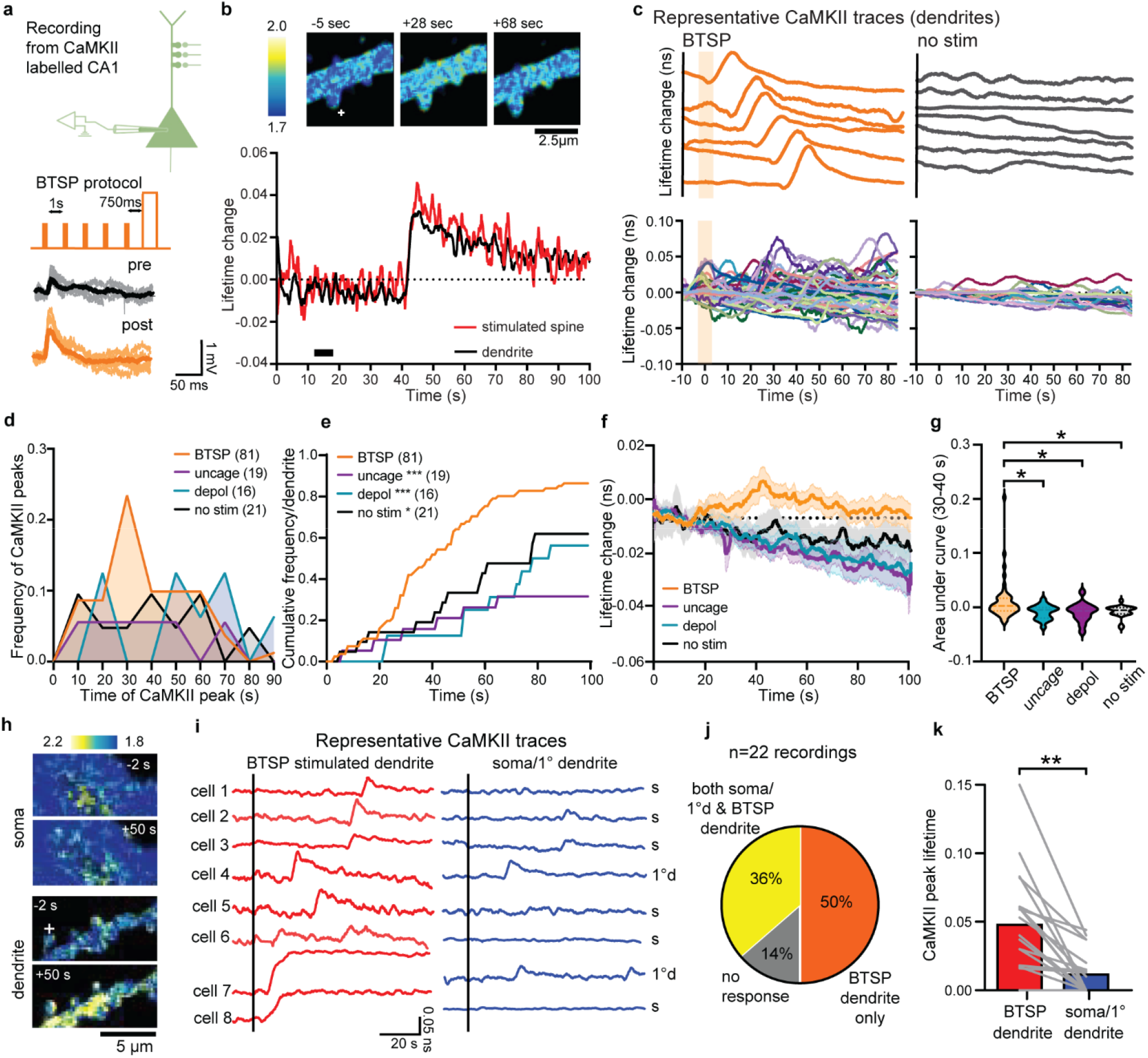
Dendritic, delayed, and stochastic CaMKII activation (DDSC) during BTSP induction. **a,** Schematics of the experimental setup (top), BTSP protocol (mid), and raw traces of 10 EPSPs and mean pre and post-BTSP protocol (bottom). **b,** A representative fluorescence lifetime images (top) and time course (bottom) of a dendritic segment expressing 2dV-Camui during BTSP induction. “+” depicts the BTSP stimulated spine. Images are acquired at 128 ms / frame and the time course was filtered with a moving average over 60 frames. **c,** Top, 6 representative traces of delayed dendritic CaMKII activity (filtered) measured as fluorescence lifetime changes of 2dV-Camui in dendrites in BTSP and non-stimulated (non-stim). Bottom, Lifetime change plots of all dendritic activity in BTSP (n=81), non-stim (n=21). The shaded bars show the BTSP protocol. **d,** Frequency of DDSC onsets after BTSP induction (orange) compared with uncaging only (uncage), current injection only (depol), and no stimulation (no-stim) controls. Numbers of dendrites are in the figure. **e,** Cumulative frequency of **d**. *p < 0.05, ***p < 0.001, Kolmogorov-Smirnov test against BTSP. **f-g,** Average time course (**f**) and area under the curve (30-40 s post-BTSP) (**g**) of mean lifetime change of CaMKII activity under BTSP, non-stimulated (no-stim), current injection only (depol), and uncaging-only (uncage) conditions. *p<0.05, one-way ANOVA followed by multiple comparison tests with Dunnett’s correction). **h-k,** DDSC does not propagate to the soma or primary dendrites. **h,** Representative fluorescence lifetime images of 2dV-Camui lifetime changes before (−10 sec) and after (+50 c) BTSP protocol at the secondary dendrites or the soma. **i,** Representative 2dV-Camui traces of the dendrite (red) and the soma (s) or the primary dendrite (1°d) (blue) from 8 different cells. In some samples, soma was not in the same z plane as the stimulated dendrite, and thus primary dendrite was used. **j,** Pie chart showing that out of 22 recordings, 50% showed an increase in CaMKII activity, specifically in the dendrites but not in the soma or primary dendrite. 36% of the recordings showed a CaMKII activity increase in both stimulated dendrite and soma/primary dendrite, ad 14% of the dendrites showed no detectable CaMKII activity. **k,** The amplitude of DDSC in BTSP-induced dendrites is significantly higher than that in the soma or primary dendrite. ***p<0.001, paired t-test.

To investigate the CaMKII activity during BTSP, we performed a whole-cell patch-clamp electrophysiology on 2dV-Camuiα expressing CA1 neurons in organotypic slice culture and imaged CaMKII activity during BTSP induction (**Fig. 3a**). While there was no CaMKII activity during BTSP induction (**Fig. 3b-d**), we observed CaMKII activation both in stimulated spines and adjacent shafts with a delay of tens of seconds (**Fig. 3b**, **c**). This delayed global CaMKII activity shows stochasticity in its timing. We found 76% of dendrites show delayed CaMKII activation, with peak timings ranging from 0 to 100 s, particularly clustered around 30-40 s (**Fig. 3c-e**, n=81 dendrites). In control neurons without any stimulation, we also observed CaMKII activation, but with a significantly lower frequency (**Fig 3c-e**, no-stim, n=21). Similarly, uncaging pulses without current injection (**Extended Fig. 3-1** uncaging, n=19), and current injection without uncaging (**Extended Fig. 3-1**, depolarization, n=16) show CaMKII activation with a frequency substantially lower than in BTSP-induced dendrites. In a subset of experiments where we recorded CaMKII activation before BTSP induction for a longer time (100 s) as well as after BTSP, and we also observed a significant increase in CaMKII activation after BTSP induction (**Extended Fig. 3-1**). The average time course (**Fig. 3f**) showed a downward drift in the measurement, perhaps due to photo-bleaching. However, on top of the drift, there exists a substantial elevation of CaMKII activity in BTSP-induced dendrites ∼30-40 secs after BTSP protocol, but not in the control groups. The area under the curve in BTSP-dendrites from 30-40 seconds showed significantly higher CaMKII activation compared with the three controls (**Fig. 3g**, two-way ANOVA with Tukey’s correction). We found no difference in the peak amplitude of CaMKII activity between BTSP-induced and control conditions (**Extended Fig. 3-1**), indicating that BTSP increases the frequency, but not the amplitude, of CaMKII activation.

In a subset of the above experiments, we recorded EPSP in stimulated spines and found similar BTSP potentiation in the CaMKII labeled neurons (**Extended Fig. 3-2a**). Moreover, there is an inverse correlation between the magnitude of potentiation and the time of CaMKII occurrence after BTSP induction (**Extended Fig. 3-2b**), suggesting that earlier CaMKII activity tends to result in a higher magnitude of potentiation. We also performed CaMKII imaging experiments during the cBTSP protocol (**Extended Fig. 3-3**). Similar to forward BTSP, we did not observe any CaMKII during cBTSP protocol but found a delayed stochastic CaMKII activation at 30-60 sec after induction of cBTSP (**Extended Fig. 3-3**). The peak amplitude and the frequency of CaMKII activity were similar between BTSP and cBTSP (**Extended Fig. 3-3**). These experiments suggest that BTSP or cBTSP protocol induces dendritic, delayed, and stochastic CaMKII activation (DDSC). DDSC requires both pre- and postsynaptic components, suggesting that CaMKII upstream signaling can integrate these components.

To investigate the spatial profile of DDSC, we imaged CaMKII activity simultaneously in stimulated dendrites and the soma or primary dendrite following BTSP induction (**Fig. 3h**). We found that DDSC is more predominant in the BTSP-induced dendrites than in the soma or primary dendrites (**Fig. 3i-k**). We found that 50% of the recordings show CaMKII activity specific to the BTSP-stimulated dendrites, while 36% of them show CaMKII activity both in the BTSP dendrite and the soma (or primary dendrite) (**Fig. 3j**). Furthermore, the peak CaMKII amplitude in dendrites was significantly higher in the BTSP-induced dendrites than in the soma or primary dendrites (**Fig. 3k**). We found similar dendrite predominant CaMKII activity in cBTSP protocol as well (n=13, **Extended Fig. 3-4a**), where 48% of the recordings showed dendritic CaMKII only in the stimulated dendrites and showed a significantly larger CaMKII peak amplitude in stimulated dendrites compared to somatic CaMKII (**Extended Fig. 3-4b**). Overall, these experiments suggest that DDSC is compartmentalized to the stimulated dendrite and does not spread throughout the cell.

**Figure 4:**
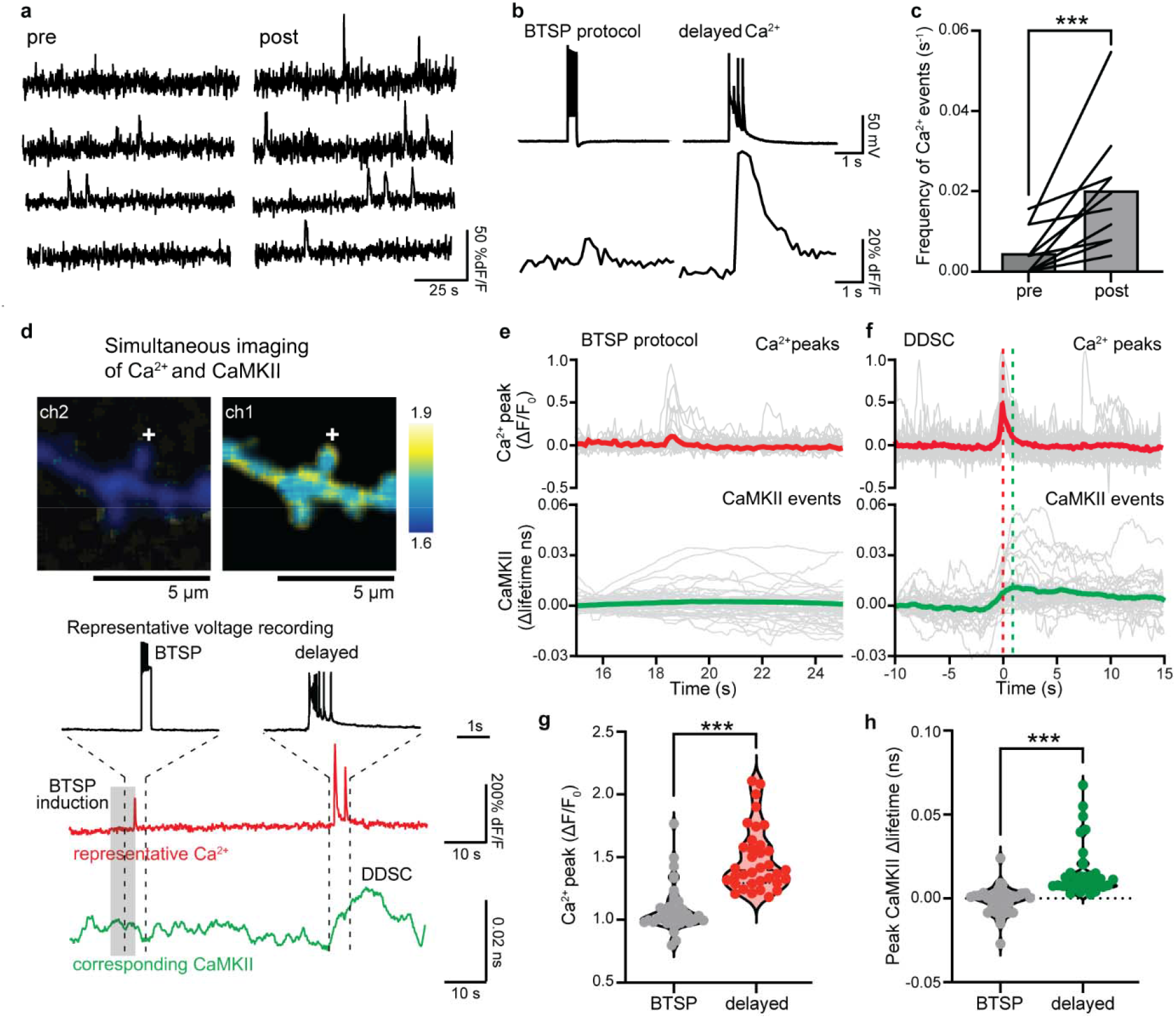
Ca^2+^ imaging shows an increase in Ca^2+^ spikes after BTSP. **a,** Snippet of Ca^2+^ traces before (4 min) and after (4 min) BTSP induction in neurons filled with Cal-590 (50 µM) through patch pipette. **b,** Representative traces of voltage recordings and corresponding Ca^2+^. **c,** The frequency of Ca^2+^ events before after BTSP induction (n = 12). Paired two-tailed t-test, *p<0.05. **d,** Top: Representative fluorescence lifetime images of a dendrite during simultaneous Ca^2+^ (Cal-590) and CaMKII imaging. Bottom: Representative voltage, Ca^2+^ and CaMKII time courses. **e,** Average Ca^2+^ elevation (red) and CaMKII activity (green) in the stimulated dendrite during BTSP protocol (n = 37). There was no detectable CaMKII activation during the BTSP protocol. **f,** Average of Ca^2+^-event-triggered average of Ca^2+^ (red) and CaMKII events (green) following BTSP induction (n = 37). **g-h,** Summary of Ca^2+^ (**g**) and CaMKII (**h**) peak amplitudes observed during BTSP protocol (as in **e**) and after BTSP induction (as in **f**). ***p < 0.001 two-tailed unpaired test.

### CaMKII activity is associated with Ca^2+^ and plateau potentials in dendrites

Since CaMKII activation requires Ca^2+^ elevation ^11^, we hypothesized that DDSC is associated with dendritic Ca^2+^ elevation. To test this hypothesis, we performed Ca^2+^ imaging by filling the cell with Cal 590 (50-100 μM) dyes in a whole-cell configuration for 4 min before and after the BTSP protocol (**Fig. 4a-c**). We confirmed that applying the BTSP protocol to these neurons potentiates EPSPs (**Extended Fig. 4-1**). Consistent with DDSC, we found Ca^2+^ elevations in close correlation with plateau potentials during 4 min recordings after application of the BTSP protocol (**Fig. 4b**). The frequency of these delayed Ca^2+^ peaked around 20-30 s, again consistent with DDSC (**Extended Fig. 4-2**). Like DDSC, Ca^2+^ elevations also occur before stimulation but are significantly less frequent (**Fig 4c**). The majority of Ca^2+^ events were correlated with plateau potentials (Extended Fig. 4-2)

To examine whether these delayed Ca^2+^ transients and the plateau potential correspond to CaMKII activity, we performed simultaneous Ca^2+^ and CaMKII imaging by filling 2dV-Camui transfected CA1 neurons with a Ca^2+^ indicator Cal-590 (50 μM) through patch pipette (**Fig. 4d**). Ca^2+^ elevations during the BTSP protocol, likely due to backpropagating action potentials, were much smaller in amplitude and did not show associated CaMKII events (**Fig. 4e**). However, after BTSP, we observed CaMKII activity, corresponding to DDSC, associated with Ca^2+^ elevation and the plateau potential (**Fig. 4d**, **Extended Fig. 4-1d**). Ca^2+^ triggered average of CaMKII activity clearly showed that the onset of CaMKII activation is temporally aligned with Ca^2+^ elevation (**Fig. 4e).** Both Ca^2+^ and CaMKII events are much larger during delayed events compared to those during the BTSP protocol (**Fig. 4g**, **h**).

### DDSC is required and sufficient as an instructive signal for BTSP

To test whether DDSC is essential in BTSP induction, we transduced neurons with a photo-inducible CaMKII inhibitor paAIP2 ^22^ using AAV, and inhibited CaMKII 0 or 30 s after BTSP induction by illuminating with blue light (470 nm, BL) (**Fig 5a, b**). Control CA1 neurons, which express paAIP2 but with no BL exposure, showed potentiation in EPSP after the BTSP protocol (89 ± 20%, n = 12) (**Fig. 5c**, **d**). However, when we inhibited CaMKII 0 or 30 s after BTSP protocol by BL, significantly less synaptic potentiation was induced (11 ± 14% for BL with 0 s delay, n = 10, 14 ± 8% for 30 s delay, n = 11, **Fig. 5c-g**). These results suggest that DDSC is necessary for BTSP induction.

**Figure 5:**
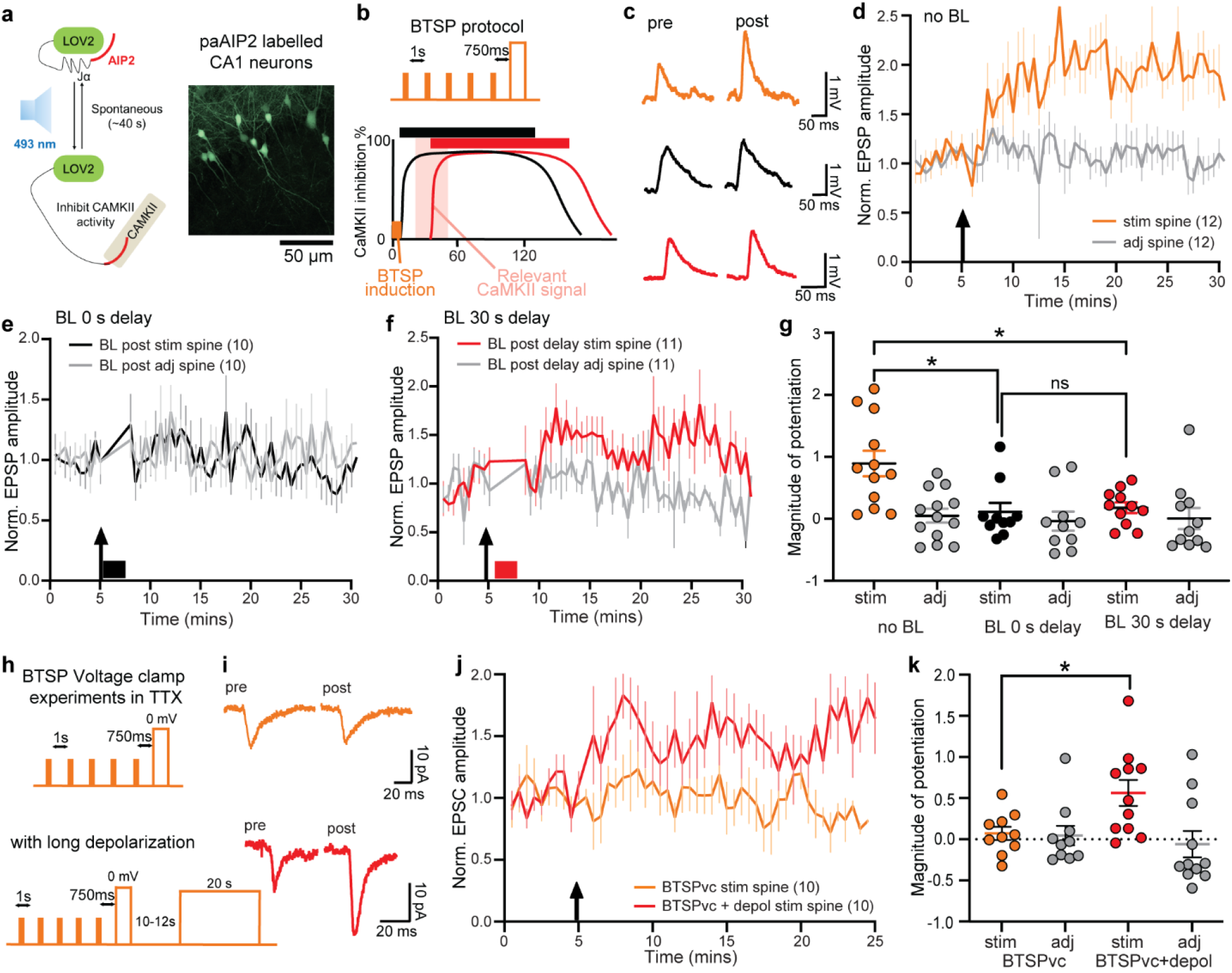
Optical inhibition and voltage clamp experiments show that DDSC plays a critical instructive role in BTSP. **a,** Schematics of photoactivatable CaMKII inhibitor paAIP2. Left: Schematics of paAIP2. Upon absorption of blue light (BL, 470 nm), light-oxygen voltage 2 (LOV2) domain changes its conformation, and exposed AIP2 inhibits CaMKII. When the BL is stopped, the paAIP2 becomes inactive within ∼40 s. Right: paAIP2-P2A-EGFP labeled CA1 neurons. **b,** Schematics of the two separate CaMKII inhibition experiments. CaMKII was inhibited for 2 mins, 0 (black) sec or 30 s (red) after the BTSP protocol (orange). **c,** Representative EPSP traces of a stimulated and adjacent spine (average of 10 traces) pre and post BTSP induction in paAIP2 labeled neurons where no BL stimulation was given (orange), or BL with 0 s delay (black) or 30 s delay (red). **d-f**, Normalized EPSP time course of EPSPs in response to BTSP in the stimulated spines (n=12) but not in the adjacent spines (n=12) for no BL (**d**), BL with 0 s delay (n=10) (**e**) or BL with 30 s delay (n=11) (**f**). Arrow depicts the timing of BTSP induction (also in e and f). **g,** Summary of the magnitude of EPSP potentiation (25-30 min) in stimulated and adjacent spines in no BL, BL 0 s after BTSP protocol (BL 0 s delay), and BL 30 s after BTSP protocol (BL 30 s delay). *p < 0.05, Two-way ANOVA followed by posthoc multiple comparisons with Tukeys’ correction. **h,** Top: BTSP protocol in voltage clamp (BTSPvc), where a train of 5 uncaging pulses (1 Hz) was paired with depolarization to 0 mV for 300 ms with 750 ms delay after the last pulse under the presence of TTX. Bottom: A protocol to artificially induce delayed CaMKII activity in addition to BTSPvc. We applied additional depolarization ∼10-12 s after BTSPvc (BTSPvc+depol). **i,** Representative EPSP traces of stimulated spine (average of 10 traces) before (black) and 20 mins after (orange) BTSPvc or BTSPvc+depol. **j,** Normalized EPSC amplitude time course in response to BTSPvc protocol (orange) and BTSPvc plus delayed depolarization (red). Arrow is the timing of BTSPvc. Numbers of cells are in the figure. **k,** Summary of the magnitude of potentiation of stimulated and adjacent spines during BTSPvc and BTSPvc+depol (* p < 0.05, two-way ANOVA followed by posthoc multiple comparison’s test (Tukey’s correction).

To study whether DDSC is sufficient as an instructive signal in BTSP, we applied BTSP protocol under inhibition of Ca^2+^ spikes by the voltage clamp and bath-applied TTX (1 µM) and then artificially induced delayed Ca^2+^-CaMKII signaling by a long, delayed depolarization (10-12 s delay, 20 s width) ^19^ (**Fig. 5h**). In control neurons, a protocol similar to BTSP, a train of 5 uncaging pulses at 1 Hz paired with depolarization to 0 mV after a 750 ms delay, failed to induce synaptic potentiation (7 ± 8%, n=10) (Fig. **5i-k**). However, in separate experiments, when the long depolarization pulse was delivered 10-12 s after the BTSP protocol, we observed a potentiation of EPSC amplitude in the stimulated spines (56.3 ± 16%, n = 11) (**Fig. 5i-k**), suggesting that the depolarization-induced CaMKII ^19^ provided instructive signals. Taken together with optogenetic experiments, these experiments demonstrate that depolarization that likely activates global CaMKII provides an instructive signal essential for inducing synapse-specific BTSP.

### Intracellular calcium release is required for the induction of BTSP and DDSC

A previous study has shown that intracellular Ca^2+^ release from internal stores is required for in vivo BTSP-induced place cell formation ^26^. Furthermore, the intracellular store-induced Ca^2+^ release has been shown to be induced by IP3-dependent mechanisms ^27^. Thus, we investigated the role of intracellular Ca^2+^ release in BTSP and DDSC using thapsigargin (1 μM), which depletes internal stores, or xestospongin C (XestC, 1 μM), which inhibits IP3R. We found that both thapsigargin and XestC significantly inhibited BTSP-induced synaptic potentiation compared to vehicle (DMSO) (**Fig. 6a**, **b**). Thus, IP3-dependent intracellular Ca^2+^ release from internal stores is required for BTSP. Moreover, CaMKII imaging showed that DDSC was impaired in the presence of thapsigargin or XestC (**Fig 6c-f**). Both drugs reduced the frequency (**Fig. 6d**, **e**) and peak amplitude of DDSC (**Fig. 6f**). Overall, these experiments suggest that Ca^2+^ release from internal stores is required for BTSP and DDSC.

**Figure 6:**
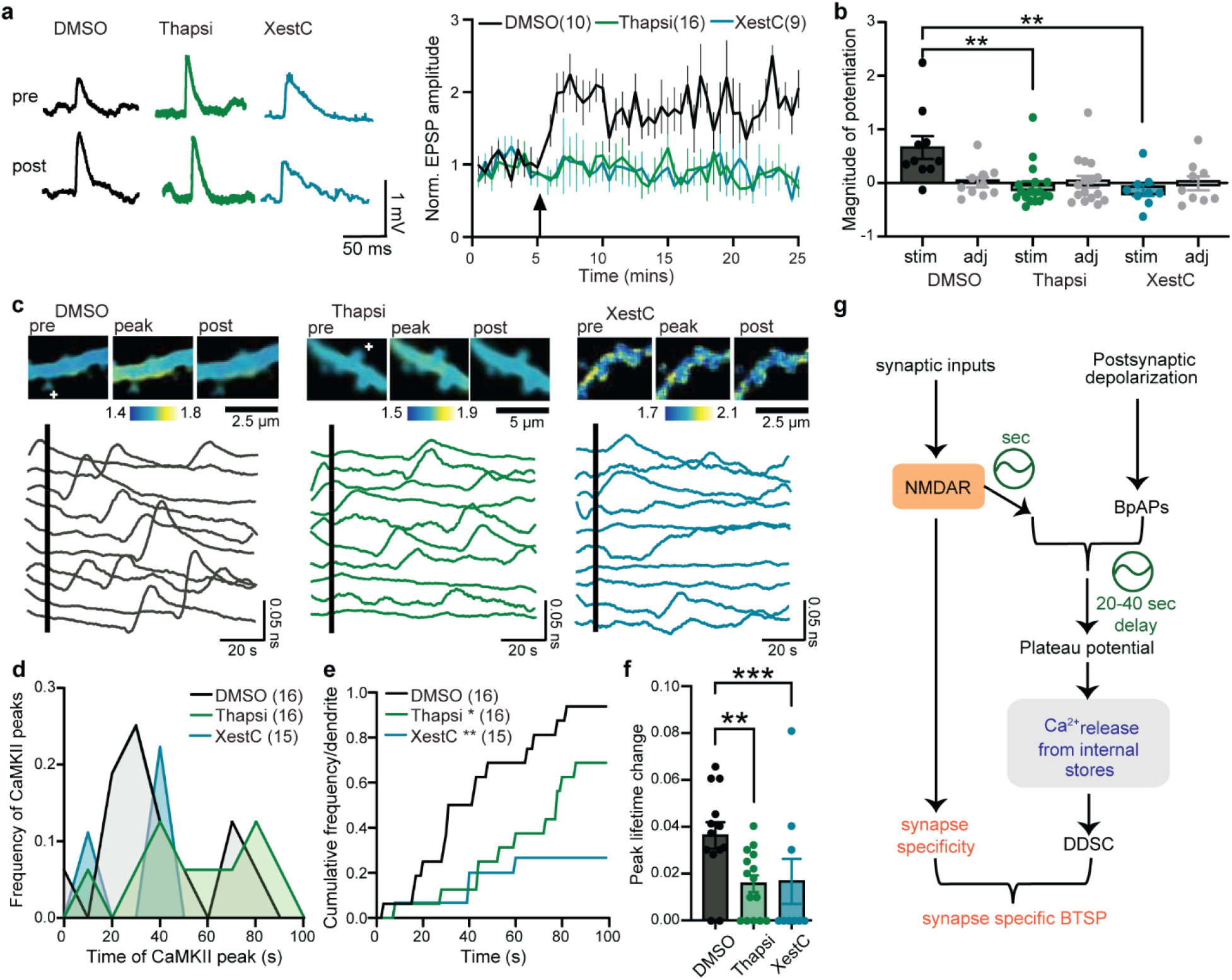
Calcium release from internal stores underlies BTSP and DDSC. **a,** BTSP-induced synaptic potentiation is impaired by depleting Ca2+ internal store with thapsigargin (Thapsi, 1 µM) or inhibiting IP3R with Xestospongin-C (XestC, 1 µM) compared with vehicle (DMSO). EPSP traces (left) and averaged synaptic potentiation in the stimulated spines (right) are shown. Numbers of samples are in the figure. **b,** Summary of the magnitude of EPSP potentiation for data in (**a**) **p < 0.01, Two-way ANOVA, with multiple comparison test (Tukey’s correction). **c,** Top: Fluorescence lifetime images of a dendrite showing lifetime change before, during and after a CaMKII peak in DMSO (left), thapsigargin (mid), and XestC (right). Bottom: 10 smoothened dendritic lifetime traces showing DDSC under each condition. **d,** The time course of the frequency of DDSC onset under DMSO, thapsigargin, or XestC conditions. **e,** Cumulative histogram of (**d**) *p <0.05, ***p<0.001 Kolmogorov-Smirnov test. **f,** Summary of peak DDSC amplitude. **p<0.01, ***p<0.001. Two-way ANOVA with multiple comparisons (Tukey’s correction). **g,** A revised model of BTSP induction. Synaptic inputs activate NMDAR-dependent signaling to prime the stimulated synapses. Combined synaptic inputs and postsynaptic activation lead to delayed plateau potentials. Ca^2+^ signal is further enhanced by intracellular Ca^2+^ release, leading to delayed CaMKII activation (DDSC). DDSC acts as an instructive signal with an extended time window of 20-40 s. Additional signals in spines must provide Synapse specificity (orange).

## Discussion

BTSP has been a leading model to explain the induction of CA1 place cells. Our uncaging-evoked behavioral time scale plasticity (BTSP) can be induced in a synapse-specific manner, like Hebbian plasticity ^28,29^. This also supports previous studies demonstrating that input-specific dendritic plasticity underlies the place cell formation at a specific location ^23,30,31^.

Although the synapse-specific role of CaMKII in synaptic potentiation has been speculated, we did not observe any detectable CaMKII activation during the BTSP protocol, even with our improved CaMKII sensor. However, we cannot rule out the possibility that there is still some activation of CaMKII below our detection limit during BTSP. Instead, we observed a dendritic, delayed, and stochastic CaMKII activity that spreads to the dendrite and nearby spines (DDSC). DDSC appears restricted to BTSP-induced dendrites and did not spread throughout the cell. Our photo-inhibition and voltage-clamp experiments suggest that DDSC plays an essential role in BTSP induction.

It is mechanistically intriguing how the pre and postsynaptic components can get paired over several hundred milliseconds and still result in an NMDAR-dependent and synapse-specific plasticity. Although the time constant of CaMKII activation during Hebbian plasticity matches with the eligibility trace of BTSP, our study suggests that CaMKII is not the eligibility trace, since CaMKII activation during BTSP neither is specific to the stimulated synapse nor active for the behavioral time scale. However, the global and delayed nature of DDSC would be consistent with its role as an instructive signal, although it provides a time window much larger than the proposed instructive signal ^32^. Since DDSC requires both pre- and postsynaptic components, additional biochemical signaling must exist upstream of CaMKII signaling (**Fig. 6g**). Furthermore, since BTSP-induced synaptic potentiation is spine-specific, the protocol needs to activate synapse-specific signaling to “prime” the stimulated spine, potentially through the metabotropic function of NMDA receptors ^33,34^. Overall, there are at least two time scales in this model, one for integrating pre- and postsynaptic inputs over the behavioral time scale (∼1 s), and the other for associating the synapse-specific priming signal and the instructive signal via plateau and DDSC (20-40 s). The integration during the behavioral time scale does not need the synapse specificity, as pre- and post-signal integration can occur in the dendrite, leading to delayed plateau potentials and DDSC (**Fig. 6g**). The signal association during the slow time scale appears to give rise to the synapse specificity of BTSP (**Fig. 6g**).

BTSP-induced plateau potentials require voltage-gated Ca^2+^ channels to elevate Ca^2+^^17^. Since our results indicate that DDSC and BTSP need IP3-dependent Ca^2+^ store release, additional amplification of Ca^2+^ by Ca^2+^-induced Ca^2+^ release (CICR) may be required for high Ca^2+^ elevation sufficient for DDSC and BTSP (**Fig. 6g**). Alternatively, IP3-induced Ca^2+^ release may be responsible for generating delayed Ca^2+^ plateau, as Ca^2+^ store release can be stochastic and global ^27^. The requirement of Ca^2+^ store release is consistent with the fact that our protocol did not induce BTSP in basal dendrites. It has been reported that basal and apical dendrites have different properties of the ER-mitochondrial coupling, which is essential for BTSP during place cell formation ^26^.

A brief current injection during our BTSP protocol did not result in a sustained plateau potential ^35^. This suggests that BTSP does not require plateau potential at the moment of the induction protocol. Instead, BTSP can facilitate the later induction of plateau potentials associated with DDSCs. This induction of plateau potentials and DDSCs by BTSP provides an extended time window of tens of seconds for associating temporarily separated events.

## Acknowledgments

The authors wish to express their gratitude to Dan Dombeck, Hidehiko Inagaki, Lesley Colgan and Krithika Ramachandran for helpful discussions on the manuscript. We would also like to acknowledge David Kloetzer for lab management, Yuki Hayano, Kathy Liu and Irena Suponitsky-Kroyter for providing their technical assistance in the paper. Lastly, we would like to thank MPFI ARC, including Elizabeth Garcia and Amanda Coldwell, for animal care and maintenance. This work was supported by National Institute of Heath grants R35NS116804 (RY), R01MH080047 (RY), U01NS128655 (RY), R01-NS-111745 (H.F), RF1-NS-113632 (H.F), and R01-MH-085926 (H.F).

## Methods

### Animals

All experimental procedures were approved and carried out in accordance with the regulations of the Max Planck Florida Institute for Neuroscience Animal Care and Use Committee as per the guidelines by the US National Institutes of Health. P4-P8 mouse pups from both sexes were used to prepare organotypic slices for imaging studies. We used *Camk2a*^T286A^ mice to test the requirement of CaMKII in BTSP experiments ^25^.

### Plasmid constructs

We fused two dimVenus (Venus_A206K, Y145W_) and mEGFP (EGFP_A206K_) to rat CaMKIIα subunit (2dV-Camuiα) ^19,36^. T286A or T305D/T306D 2dV-Camuiα mutants were constructed by restriction digestion and ligation.

### Organotypic hippocampal slice cultures and transfection

Organotypic hippocampal slices were prepared from wild-type or transgenic postnatal 4-8 day-old mouse pups of both sexes as previously described ^37^. In brief, the animal was anesthetized with isoflurane, after which it was quickly decapitated and the brain removed. The hippocampi were dissected and cut into 350 µm thick coronal hippocampal slices using a McIlwain tissue chopper (Ted Pella, Inc) and plated on hydrophilic PTFE membranes (Millicell, Millipore) fed by culture medium containing MEM medium (Life Technologies), 20% horse serum, 1 mM L-Glutamine, 1 mM CaCl_2_, 2 mM MgSO_4_, 12.9 mM D-Glucose, 5.2 mM NaHCO_3_, 30 mM HEPES, 0.075% Ascorbic Acid, 1 µg/ml insulin. The slices were incubated at 37 °C in 5% CO_2_. After 7-12 days in culture, CA1 pyramidal neurons were transfected with biolistic gene transfer using 1.0 µm gold beads (8–12 mg) coated with 2dV-Camuiα (50 μg) ^38^. Neurons expressing 2dV-Camuiα were imaged 1–5 days after transfection.

### Acute slice preparation

Male mice (P25–P35) were sedated by isoflurane inhalation and perfused intracardially with a chilled choline chloride solution. The brain was removed and placed in the same choline chloride solution composed of 124 mM Choline Chloride, 2.5 mM KCl, 26 mM NaHCO_3_, 4 mM MgCl_2_, 1.2 mM NaH2PO_4_, 10 mM Glucose, and 0.5 mM CaCl_2_, pH 7.4 equilibrated with 95%O_2_/5%CO_2_. Coronal hippocampal slices (300 μm) from both hemispheres were cut using a vibratome (V1200, Leica) and maintained in a submerged chamber in ACSF at 32°C for 1h and then at room temperature in oxygenated ACSF.

### Two-photon glutamate uncaging

Two-photon glutamate uncaging was performed during BTSP and structural LTP experiments in organotypic hippocampal cultures and acute hippocampal slices as described previously ^39^. Experiments were performed in a small recirculating volume (∼ 8 ml) of continuously oxygenated ACSF containing 4 mM 4-methoxy-7-nitroindolinyl-caged-l-glutamate (MNI-caged glutamate). Ti: Sapphire laser tuned at a wavelength of 720 nm to uncage MNI-caged glutamate in a small region ∼0.5 μm from the spine. For structural plasticity experiments, 30 uncaging pulse, 0.5 Hz train was given. The power of the laser was set to 2.7 mW measured at the objective. These structural plasticity experiments were performed in Mg^2+^ free artificial cerebral spinal fluid (ACSF; 127 mM NaCl, 2.5 mM KCl, 4 mM CaCl_2_, 25 mM NaHCO_3_, 1.25 mM NaH_2_PO_4_ and 25 mM glucose) containing 1 μM tetrodotoxin (TTX) and 4 mM MNI-caged L-glutamate aerated with 95% O_2_ and 5% CO_2_. Experiments were performed at room temperature (24–26°C).

### Electrophysiology

Whole cell patch clamp electrophysiology experiments were combined with glutamate uncaging to induce BTSP at individual dendritic spines. The cells were first visualized in a bright field, or for the labelled cells, epifluorescence microscopy. The patch pipette (with tip resistance 2-5 MΩ) included the internal solution containing 145omM K gluconate, 14omM phosphocreatine, 4 mM NaCl, 0.3omM NaGTP, 4omM MgATP, 3omM L-ascorbic acid, 50-100 µM Alexa-594, and 10omM HEPES (pH 7.4, 294 mOsm). In BTSP experiments, the EPSPs were measured under the current-clamp mode by a patch-clamp amplifier (MC-700B, Molecular Devices) and digitizer (National Instruments). After 2-5 mins of dye loading, fluorescence from Alexa-594 was used to find dendritic spines in 2pFLIM. Uncaging-evoked EPSPs were induced on 1-2 spines on a dendrite by MNI glutamate uncaging, ∼0.5 µm away from the tip of the spine. The uEPSP amplitude was 0.4-2 mV. Pairing LTP and some BTSP experiments were performed in voltage-clamp configuration, where the cells were held at −70 mV. The baseline glutamate uncaging evoked EPSC amplitude was between 5-20 pA. Some BTSP experiments were performed in voltage clamp with Cs internal solution containing 130 mM Cs-methanosulphonate, 6 mM KCl, 10 mM HEPES, 4 mM NaCl, 0.3 mM MgGTP, 4 mM MgATP, and 14 mM Tris-phosphocreatine (BTSP voltage clamp protocol). Experiments were performed at room temperature (24-26 °C). In the CaMKII imaging experiments, similar to the above experiments, Alexa 594 dye (100 µM) was loaded as a structural marker. EPSPs were measured before and after the induction of BTSP. In all whole-cell recordings, the series resistance was monitored to be between 10-40 MΩ throughout the recording.

### HeLa cell maintenance, transfection and imaging

HeLa cells (ATCC CCL-2) were grown in Dulbecco’s modified Eagle medium supplemented with 10% fetal bovine serum at 37o°C in 5% CO_2_. Plasmids were transfected into HeLa cells using Lipofectamine 3000 (Invitrogen). Imaging was performed 24-48 h following transfection in a HEPES-buffered ACSF solution (20 mM HEPES pH 7.3, 130 mM NaCl, 2 mM NaHCO_3_, 25 mM D-glucose, 2.5 mM KCl, 1.25 mM NaH_2_PO_4_) with 2 mM CaCl_2_ and 2 mM MgCl_2_ by 2pFLIM as described below. When indicated, cells were stimulated with bath application of ionomycin (1704, Tocris) and then EGTA.

### Fluorescence-coupled size-exclusion chromatography (FSEC)

Expression vector DNAs (2 µg) including Camuiα were transfected into HEK293S GnTI-cells (2 x 106 cells per well in 6-well plates) cultured in FreeStyle 293 (Thermo Fisher), using the TransIT2020 transfection reagent (Mirus Bio). Cells were harvested 48 h post-transfection, washed with ice-cold PBS, and sonicated in 250 µl of the TBS (20 mM Tris-HCl (pH 8.0) and 200 mM NaCl), using Misonix Sonicator 3000 (3 times, 30 s, power level 9.0). The lysate was ultracentrifuged at 70,000 rpm for 10 min (TLA110 rotor). The supernatant (20 µl) was loaded onto the Superose-6 size-exclusion chromatography column (10/300 GL; GE Healthcare), pre-equilibrated with TBS, and run at a flow rate of 0.4 ml/min. The eluent from the Superose-6 column was detected by a fluorometer (RF-10AXL, Shimadzu) with the following settings: excitation, 475 nm; emission, 507 nm; time increment, 0.5 s; integration time, 1 s; and recording time, 75 min. The FSEC data points were plotted by OriginPro graphic software (OriginLab).

### Fluorescence correlation spectroscopy (FCS)

HEK293FT cells (Thermo Fisher) were transfected with the plasmids using Lipofectamine 3000 (Thermo Fisher) and cultured for 2 days at 37 °C and 5 % CO_2_. After washed the plate wells once in PBS buffer, the cells were lysed for 5 min with M-PER mammalian protein extraction reagent (Thermo Scientific) including Halt protease inhibitor (Thermo Scientific) and 5 mM EDTA. The lysates were centrifuged at 20000g for 10 min and the supernatants were used for the FCS measurement by diluted 2 to 15-fold in PBS buffer including the protease inhibitor. The FCS measurements were performed at 23 °C under 2-photon microscope without laser scanning, equipped with Ti:Sapphire laser (Chameleon Ultra II, Coherent) tuned to a wavelength of 920 nm. The time-correlated single-photon counting (TCSPC) data were collected for 60-120 s using a water immersion objective (LUMPlanFL N 60× NA 1.0 W, Olympus) directly immersed in 300 µL of the lysate solution, a single-photon counting board (Time Harp 260, PicoQuant), and a software of TTTR mode real-time correlator in TimeHarp 260 v3.0. The data analysis was performed with FoCuS-point software ^40^.

### Optical CaMKII inhibition experiments

The CaMKII inhibition experiments were performed in organotypic hippocampal slices using previously described paAIP2 ^22^. In these experiments, slices were virally infected with 0.5-1 µl AAV mixture per slice (containing AAV9-Camk2a-Cre at 2 × 10^12^ vg/ml (1:1000 dilution, Addgene and rAAV8-DIO-CBA-pAAIP2-mEGP at 4.2 x 10^12^ vg/ml, UNC GTC Vector Corp) at DIV 4-6 and imaged or patched at DIV 10-13. Cells with robust EGFP expression were used for experiments. Labelled cells were patched with K glu internal and Alexa 594 dye in the patch pipette as described above. 470 nm LED light stimulation (M470L5, Thorlabs) was used to activate paAIP2.

### Two-photon microscopy and 2pFLIM

Custom-built two-photon fluorescence lifetime imaging microscope was used to perform 2pFLIM as previously described ^41^. 2pFLIM imaging was performed using a Ti-sapphire laser (Coherent, Chameleon) at a wavelength of 920 nm with a power of 1.0-1.4 mW. Fluorescence emission was collected using a water immersion objective (60×, numerical aperture 0.9, Olympus), divided with a dichroic mirror (565onm), and detected with two separated photoelectron multiplier tubes placed after wavelength filters (Chroma, 510/70-2p for green and 620/90-2p for red). Both red and green channels were fit with photoelectron multiplier tubes (PMT) having a low transfer time spread (H7422P40; Hamamatsu) to allow for fluorescence lifetime imaging. Photon counting for fluorescence lifetime imaging was performed using a time-correlated single-photon counting board (Time-harp 260, Pico-Quant) using custom software developed in C# (https://github.com/ryoheiyasuda/FLIMage_public). 2pFLIM images were collected at 64×64 pixels at the frame rate of 7.8 Hz. A second Ti-sapphire laser tuned at a wavelength of 720 nm was used to uncage MNI-caged glutamate.

### Ca^2+^ imaging

Ca^2+^ imaging was performed by loading calcium dyes Cal 590 (50 μM, AAT Bioquest) together with a structural marker Alexa 488 (100 μM, Thermo Fisher Scientific). The intensity of Ca^2+^ sensor were collected at 64×64 pixels at the frame rate of 7.8 Hz with 2pFLIM (lifetime information was not used). The Ca^2+^ response was calculated by normalizing the intensity with the intensity Alexa 488. The membrane voltage was also recorded during Ca^2+^ imaging under the current-clamp mode. In a subset of experiments, uncaging-evoked EPSPs were measured before and after BTSP induction. In experiments where simultaneous Ca^2+^ and CaMKII experiments were performed, Ca^2+^ was normalized to the average of first 100 frames before the induction of BTSP. The Ca^2+^ events were detected using a custom python code, where 3 times the standard deviation of the baseline noise was used as a detection threshold after the subtraction of basal trend line obtained by linear regression.

### 2pFLIM analysis

2pFLIM analysis was performed as previously described ^42^. To measure the fraction of the donor that was undergoing FRET with the acceptor (binding fraction), we fit a fluorescence lifetime curve summing all pixels over a whole image with a double exponential function convolved with the Gaussian pulse response function:

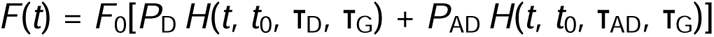

where T_AD_ is the fluorescence lifetime of the donor bound with the acceptor, *P*_D_ and *P*_AD_ are the fraction of free donor and donor undergoing FRET with the acceptor, respectively, and *H*(*t*) is a fluorescence lifetime curve with a single exponential function convolved with the Gaussian pulse response function:

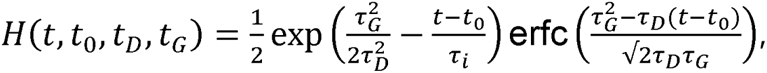

in which T_D_ is the fluorescence lifetime of the free donor, τ_G_ is the width of the Gaussian pulse response function, *F*_0_ is the peak fluorescence before convolution and *t*_0_ is the time offset, and erfc is the complementary error function.

We fixed T_D_ to the fluorescence lifetime obtained from free mEGFP (2.6 ns). For experimental data, we fixed T_D_ and T_AD_ to these values to obtain stable fitting. To generate the fluorescence lifetime image, we calculated the mean photon arrival time, <*t*>, in each pixel as:

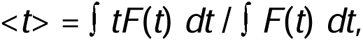

Then, the mean photon arrival time is related to the mean fluorescence lifetime, <T>, by an offset arrival time, *t_o_*, which is obtained by fitting the whole image:

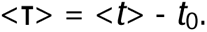

For small regions-of-interest (ROIs) in an image (spines or dendrites), we calculated the binding fraction (P_AD_) as:

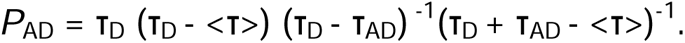

To measure the CaMKII time of occurrence and peak lifetime change in BTSP and control experiments, the raw traces were first normalized using the first 100 frames as baseline and then the normalized data was smoothened using a moving average of 60 data points. Following this processing, the time of CaMKII peak and amplitude was manually calculated on individual CaMKII traces.

### Experimental design and statistical analysis

All values are presented as mean ± SEM unless otherwise noted. Number of independent measurements/cells (n) is indicated in figures or figure legends. For our pharmacology experiments, 1-2 control experiments were performed on the same slices before the specific drug was added in the ACSF. In experiments where DMSO was used as a vehicle, we performed control experiments on different days but on the slices made from the same litter as used in the experiments. Unpaired two-tailed student’s t test was used for comparing two independent samples. One way ANOVA followed by multiple comparison tests was used for comparing more than two independent samples. Two way ANOVA followed by multiple comparison test was used to compare grouped data sets. Correlation analysis was done by computing Pearson correlation coefficients. Data smoothening, statistical tests and p values are noted in each figure legend and were computed using GraphPad Prism 7.03 for Windows, GraphPad Software, La Jolla California USA, www.graphpad.com.

## Supplementary note: Characterization of new CaMKII sensors

### Sensor Screening in HeLa cells

Original Green-Camuiα sensor (1dV-Camui) is made of CaMKIIα subunit fused with dimVenus (Venus_A206K, Y145W_ or dV) and mEGFP (EGFP_A206K_) at N- and C-termini, respectively. To improve the sensor, we screened several modified sensors (**Extended Fig. 2-1**) ^19^. We tested Camui with multiple acceptors (2dV-Camui and 3dV-Camui). Also, we tested a variant in which dimVenus is replaced by ShadowY (ShY) ^22^, which has an even lower quantum yield compared to dimVenus (1ShY-Camui and 2ShY-Camui).

In HeLa cells, we measured fluorescence lifetime changes in response to the bath application of ionomycin (3 µM) using 2pFLIM. Then, the decay kinetics of the sensors was measured by applying EGTA (8 mM). We found that all sensors show qualitatively similar kinetics (**Extended Fig. 2-1b,c**). For dimVenus, we found a larger number of accepters provide a better signal and lower basal fluorescence lifetime, due to high FRET efficiency (**Extended Fig. 2-1d, e**). However, Camui with ShY did not show a clear correlation between the number of accepters and FRET signals (**Extended Fig 2-1d, e**).

While 3dV-Camui provided the best signal among these sensors, we decided not to use this sensor because it did not express well in neurons. Among others, we found that 2dV-Camui showed the best signal-to-background ratio (**Extended Fig. 2-1b,c**), and thus further analyzed this sensor.

To validate 2dV-Camui sensor, we analyzed their mutants (**Extended Fig. 2-2**). When we mutated an autophosphorylation site that prolongs CaMKII activation (T286A) ^11,21,43–45^, the sensor signal was substantially reduced (**Extended Fig. 2-2**). Furthermore, when we eliminated calmodulin binding required for CaMKII activation (T305D and T306D) ^46,47^, the sensor signal was abolished (**Extended Fig. 2-2**). These results are consistent with previously described CaMKII behavior ^11^.

### Fluctuation correlation analysis

CaMKII holoenzyme consists of dodecamer. Thus, we next studied if they can form dodecamer using fluctuation correlation spectroscopy (FCS) of the lysate of HEK293FT cells expressing CaMKII sensors (**Extended Fig. 2-3**). FCS allows us to measure the diffusion speed of fluorescence molecules by measuring the time correlation of the fluorescence in a focused laser spot. Since the diffusion is sensitive to the hydrodynamic diameter, we can estimate the size of fluorescence particles in the lysate^48^

We found that all EGFP-CaMKII, 1dV-Camuiα, 2dV-Camui, 3dV-Camui, 1ShY-Camui, 2ShY-Camui showed similar diffusion constant (∼10 µm^2^/s), suggesting that increasing the number of accepters do not affect the hydrodynamic diameter (**Extended Fig. 2-3a**). In comparison, the diffusion constant of EGFP (84 ± 2.9 µm^2^/s) and monomeric CaMKIIα (truncated CaMKIIα[1-306]) (62 ± 2.0 µm^2^/s) was much higher consistent with their smaller hydrodynamic diameters ^48^

The diffusion speed of the sensors is somewhat smaller than previously measured diffusion of non-labeled CaMKII by dynamic light scattering (25 µm^2^/s) ^49^, perhaps because of the fluorophore fusion. To further examine if they can co-polymerize with non-labeled CaMKII, we co-expressed non-labeled CaMKII together with CaMKII sensors with different ratios. We found that, as the ratio of non-labeled CaMKII increases, the diffusion constant becomes higher, suggesting that they can copolymerize (**Extended Fig. 2-3a, b**). Also, the y-intercept, which denotes the diffusion of non-labeled CaMKII, shows the diffusion constant similar to the one previously measured.

Thus, overall, these experiments suggest that the fusion of multiple accepters do not affect the hydrodynamic radius of the molecule, suggesting that they can form normal size of holoenzyme. Also, the new CaMKII sensor (2dV-Camui) can copolymerize with non-labeled CaMKIIα.

### Fluorescence coupled size-exclusion chromatography assay

Next, we performed fluorescence coupled size-exclusion chromatography (FSEC) to measure the approximate size of the holoenzyme. Before testing the new sensors, we examined Green Camuiα, and EGFP-tagged CaMKIIα. As reported before ^19,36^, Green Camuiα shows similar peak retention time with EGFP-tagged CaMKIIα, suggesting that Green Camuiα is a dodecamer ^36^ (**Extended Fig. 2-4**). Notably, when we tested Green Camuiβ, which uses CaMKIIβ instead of CaMKIIα, we observed an additional peak corresponding to smaller complex (**Extended Fig. 2-4).** When we ran 2dV-Camui on the column, we found that this sensor showed a single peak time similar to EGFP-CaMKIIα and 1dV-Camuiα. Thus, taken together with FCS analysis, we concluded that 2dV-Camuiα can form dodecamer holoenzyme.

### Characterization of 2dV-Camui with 2-photon glutamate uncaging

Finally, we tested 2dV-Camui in dendritic spines of pyramidal CA1 neurons in organotypic hippocampal slices. We applied 30 pulses of 2-photon glutamate uncaging in the absence of extracellular Mg^2+^. Consistent with previous study ^21^, we found that 2dV-Camui is activated in a step-wise manner in response to each uncaging pulse in the stimulated spine, and then decayed with a fast time constant of 7.3 s. The decay time constant was similar to that of 1dV-Camui (7.9 s). These kinetics are consistent with our previous study using the original Green-Camuiα (1dV-Camui) ^21^. Furthermore, 2dV-Camui with T286A mutation showed much faster decay (0.74 s) (**Fig. 2**), and failed to integrate uncaging pulses (**Fig. 2**), again consistent with our previous work ^21^. Finally, we found that the sensor with T305D and T306D mutations abolished the response, as expected ^50^. Overall, our results indicate that 2dV-Camui provides signals with the kinetics similar to the original Green-Camuiα, but with ∼2 fold higher sensitivity.

---

**Extended Figure 1:**
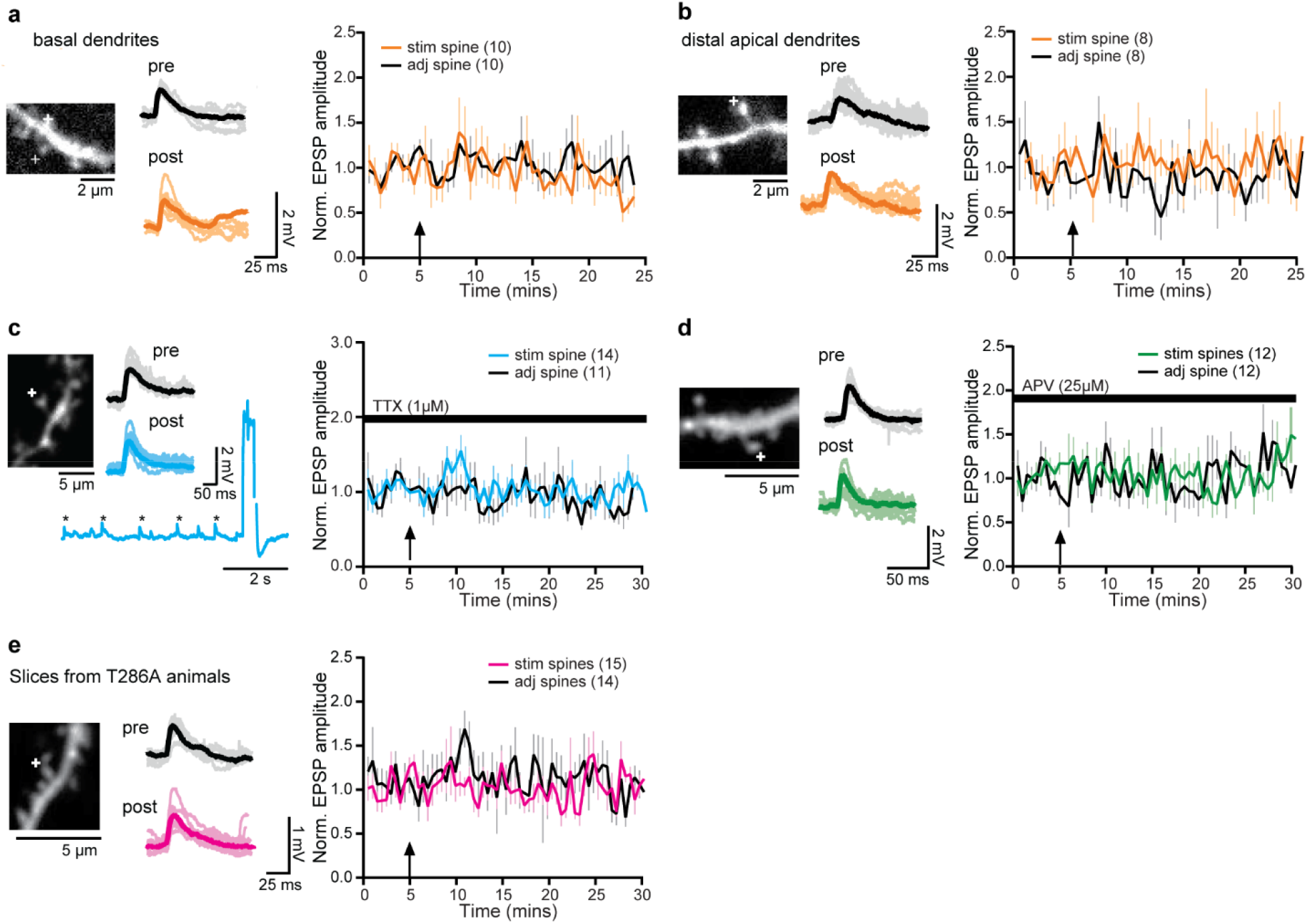
BTSP fail to induce in basal or distal synapses and is dependent on TTX, APV and CaMKII. **a,** Right: Representative basal dendritic image where BTSP was induced in one spine (+); Bottom: raw traces of EPSPs in stimulated spine before and after BTSP induction. Left: Averaged time course of normalized EPSP amplitude following BTSP induction in basal dendrites in stimulated and adjacent spines. N is indicated in the figure. **b,** Same as (**a**) but in distal dendrites. **c,** Right: Representative image of a dendritic shaft, where BTSP protocol was induced in one spine (+) in the presence of TTX (1 µM). All experiments were performed after incubating the slices for ∼30 mins in TTX. Bottom: Raw traces of EPSP in the stimulated spine in TTX. 10 EPSPs and mean before (black) and after (blue) BTSP induction. Left: Averaged time course of normalized EPSP amplitude in TTX in stimulated and adjacent spines. **d,** Right: Same as (**c**), but performed after incubating the slices for 30 mins in the presence of APV (25 µM). **e,** Right: Same as (**c**), but in hippocampal slices made from CAMKIIα^T286A^ mice.

**Extended Figure 2-1:**
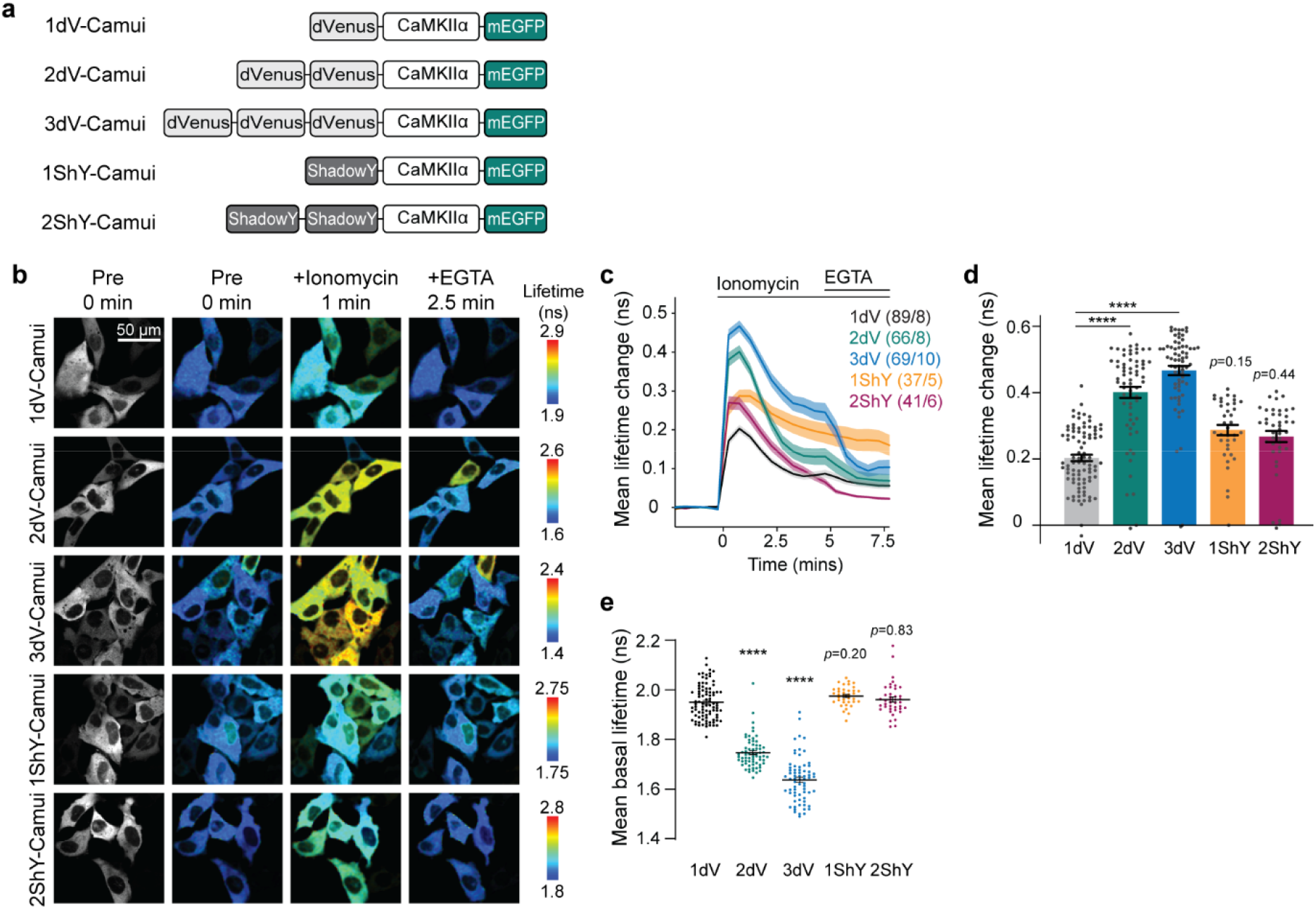
Characterization of novel conformational CaMKII FRET sensor. **a,** A schematic of screened CaMKII sensors where N- and C-termini of CaMKIIα are labeled with 1 to 3 dimVenus or 1 to 2 ShadowY acceptor(s) and donor EGFP fluorophores. **b,** Grayscale and fluorescence lifetime images of Hela cells expressing each Camuiα sensor variants. Each image is before (pre), 1 minute after ionomycin (3 µM) and 2.5 minutes after EGTA (8 mM) application. **c-d,** Averaged time course (**c**) and summary of the peak (1 min post-ionomycin application) (**d**) of fluorescent lifetime changes of 2dV-Camui (2dV) and Green-Camui (1dV) in response to bath application of ionomycin in HeLa cells. The numbers of each sample are shown in the figure (cells/cultures). **e,** Averaged basal lifetime of each CaMKII sensor. Multiplying dimVenus acceptors showed significantly lower basal lifetimes indicating higher FRET efficiency. The data are presented as meanL±LSEM., **** p < 0.0001, One-way ANOVA followed by Dunnet’s post hoc test.

**Extended Figure 2-2:**
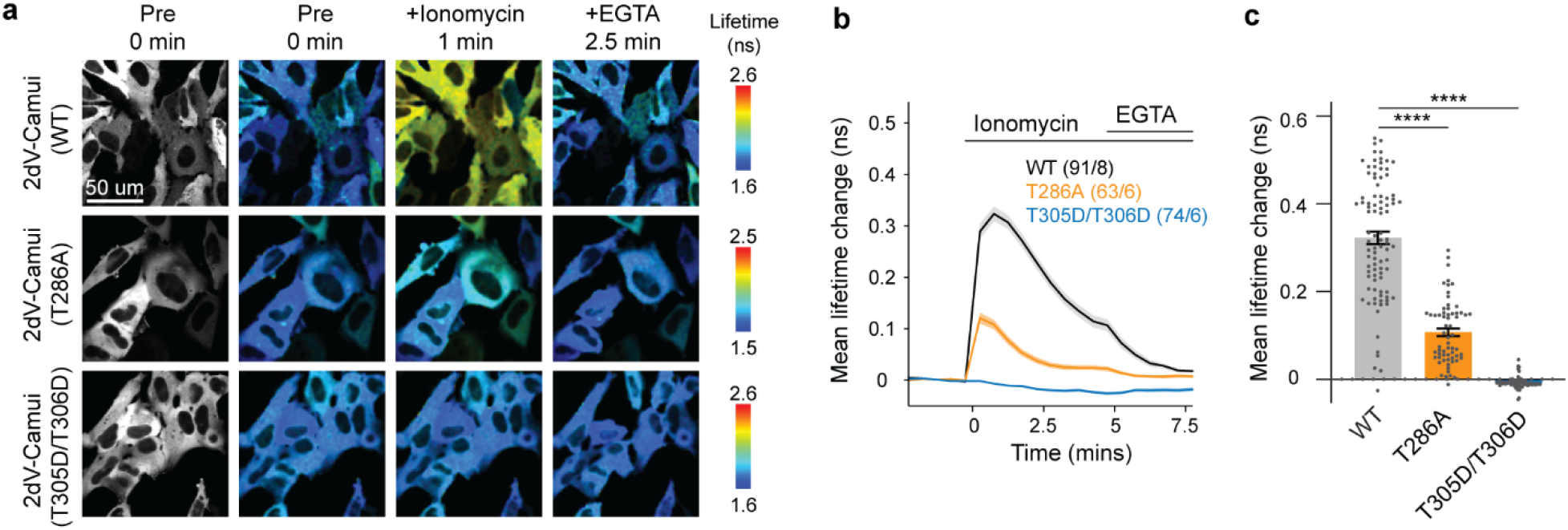
Validation of 2dV-Camui conformational CaMKII FRET sensor with wildtype, T286A or T305D/T306D CaMKII mutants. **a,** Grayscale and fluorescence lifetime images of Hela cells expressing 2dV-Camuiα sensors, which have wildtype CaMKII (WT), CaMKII mutant deficient autonomous activation (T286A), or CaMKII mutant deficient in Ca2+/calmodulin binding (T305/T306D). Each image is before (pre), 1 minute after ionomycin (3 µM) and 2.5 minutes after EGTA (8 mM) application. **b-c,** Averaged time course (**b**) and summary of the peak (1 min post-ionomycin application) (**c**) of fluorescent lifetime changes of 2dV-Camui (2dV) and Green-Camui (1dV) in response to bath application of ionomycin in HeLa cells. The numbers of each sample are shown in the figure (cells/cultures). The data are presented as meanL±LSEM., **** p < 0.0001, One-way ANOVA followed by Dunnet’s post hoc test.

**Extended Figure 2-3.**
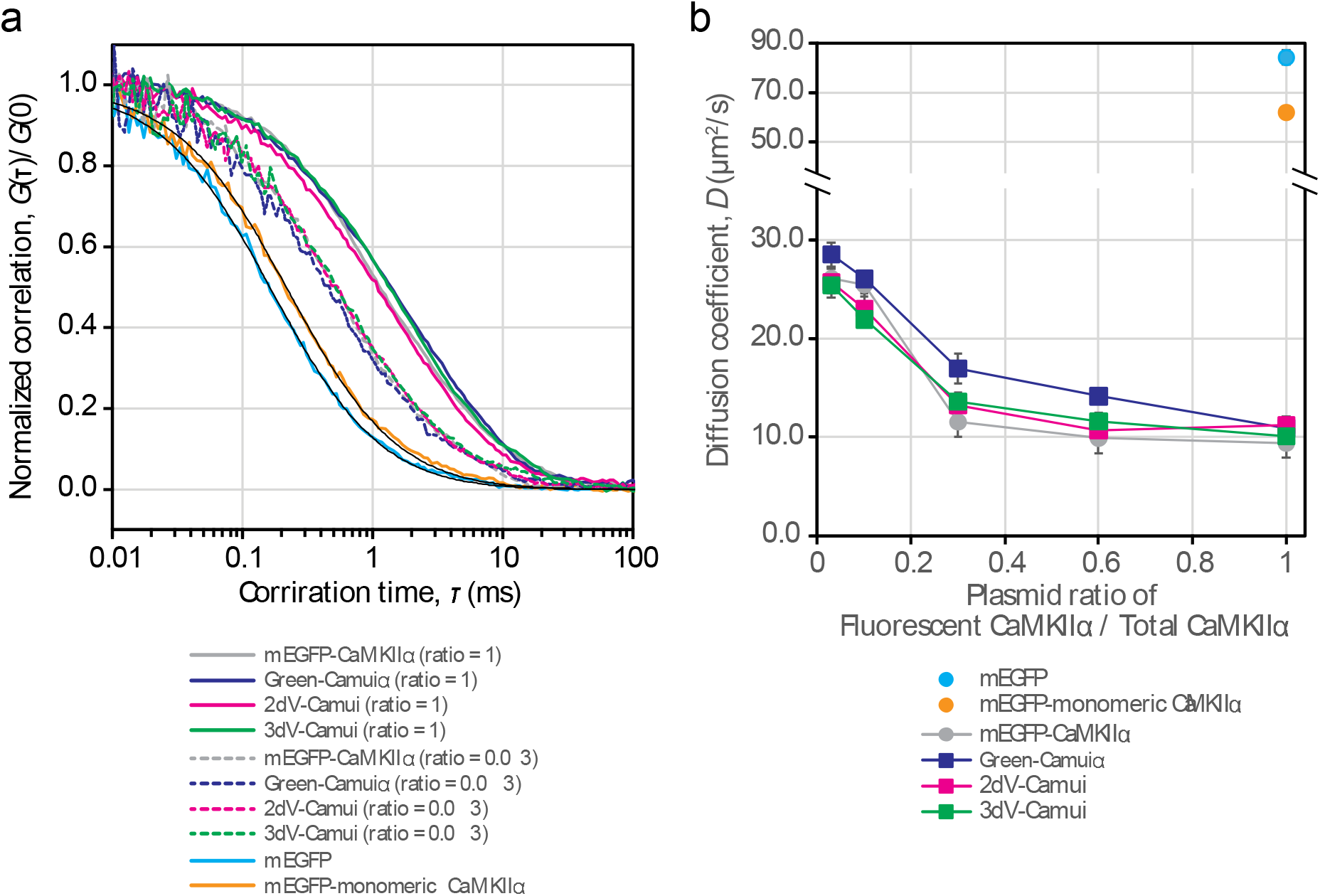
Fluctuation correlation spectroscopy (FCS) analysis of CaMKII sensors. **a,** Normalised correlation curves obtained from 2-photon TCSPC (Time-Correlated Single Photon Counting) FCS for the indicating samples in HEK293FT cell lysate. To investigate whether the sensors were co-polymerised with nonlabelled CaMKIIα, the cells were co-transfected with nonlabelled CaMKIIα at the indicating plasmid ratios. Black lines show fit curves of a correlation function given by: 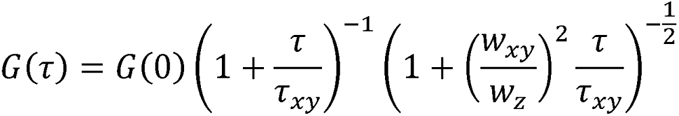 where *G*(0) is the correlation at time 0, *τ* is the correlation time, *w*_xy_ is the lateral and *w*_z_ the axial 1/*e*^2^-radii of the 2-photon excitation volume, which were measured as 0.34 µm and 1.49 µm, respectively, by scanning 0.1-µm fluorescent bead. The diffusion coefficient 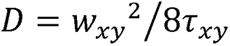 was determined from the average lateral diffusion time *τ*_xy_ obtained by curve fitting. **b,** Diffusion constants of the sensors or mEGFP-CaMKIIα co-expressed with non-labelled CaMKIIα at various plasmid ratios. The diffusion coefficients (ratio = 1) were 9.4 ± 1.5 µm^2^/s for mEGFP-CaMKIIα, 11 ± 0.6 µm^2^/s for Green-Camuiα, 11 ± 0.9 µm^2^/s for 2dV-Camui, 10 ± 0.9 µm^2^/s for 3dV-Camui, 84 μm^2^/s for EGFP, and 63 μm^2^/s for monomeric CaMKIIα (truncated CaMKIIα [1-306]), respectively. Error bars denote SD.

**Extended Figure 2-4.**
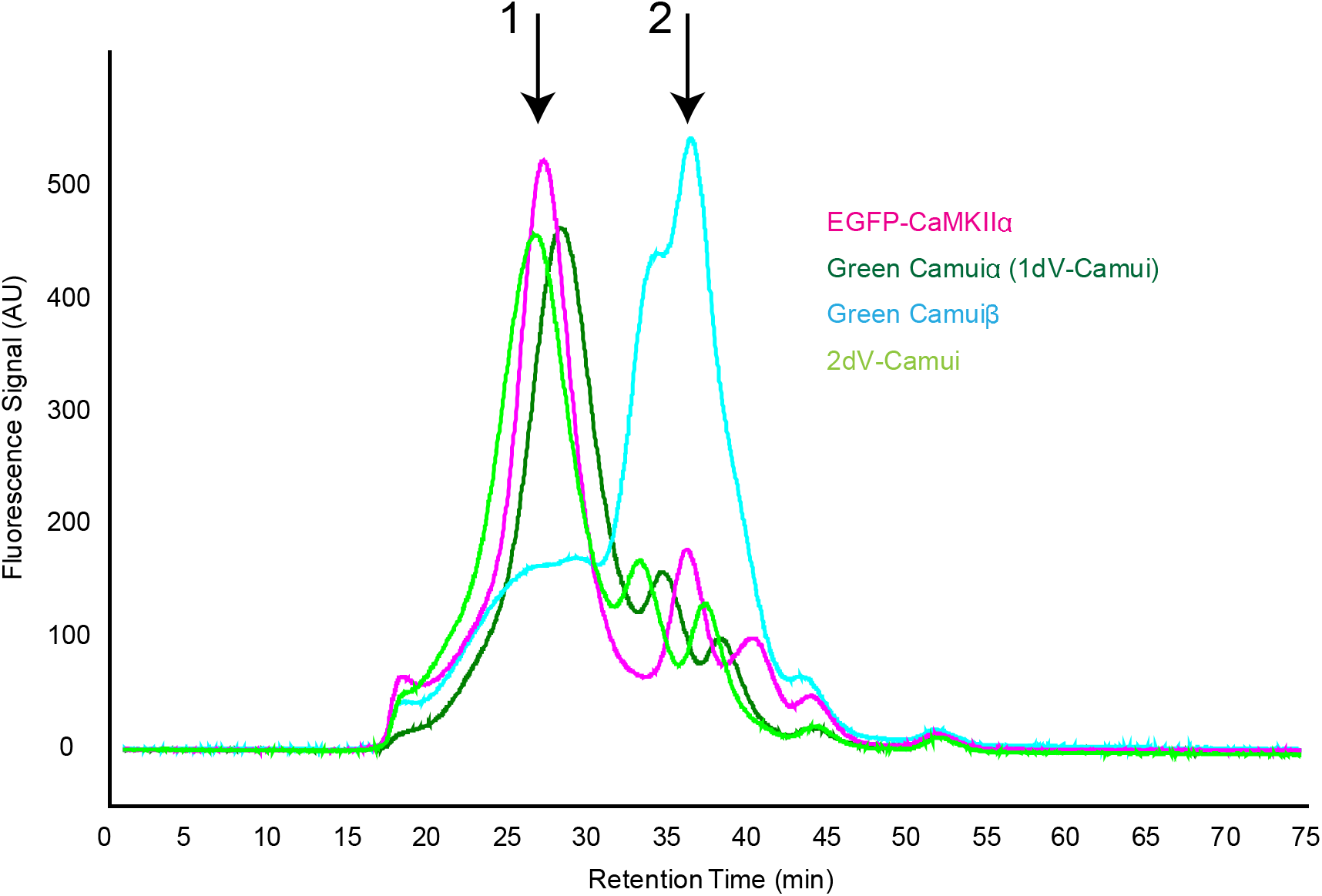
Chromatography assay of CaMKII sensors. To estimate the approximate oligomeric state of CaMKII sensors, we performed FSEC on EGFP-CaMKIIα and various Camui proteins expressed in HEK293 cells by transient transfection. The cell lysates expressing these proteins were directly injected to a Superose-6 column, and EGFP fluorescence was detected using 475/507 nm excitation/emission wavelengths. Green Camuiα (1dV-Camui), Green Camuiβ, in which CaMKIIβ subunit is used instead of CaMKIIα, and 2dV-Camui. 2dV-Camui-2dV showed a similar retention time to EGFP-CaMKIIα and 1dV-Camui (peak at ∼27 min; arrow 1), suggesting formation of a dodecamer. However, Green Camuiβ showed a high fraction of lower oligomeric species represented by a peak at ∼35 min (arrow 2).

**Extended Figure 3-1:**
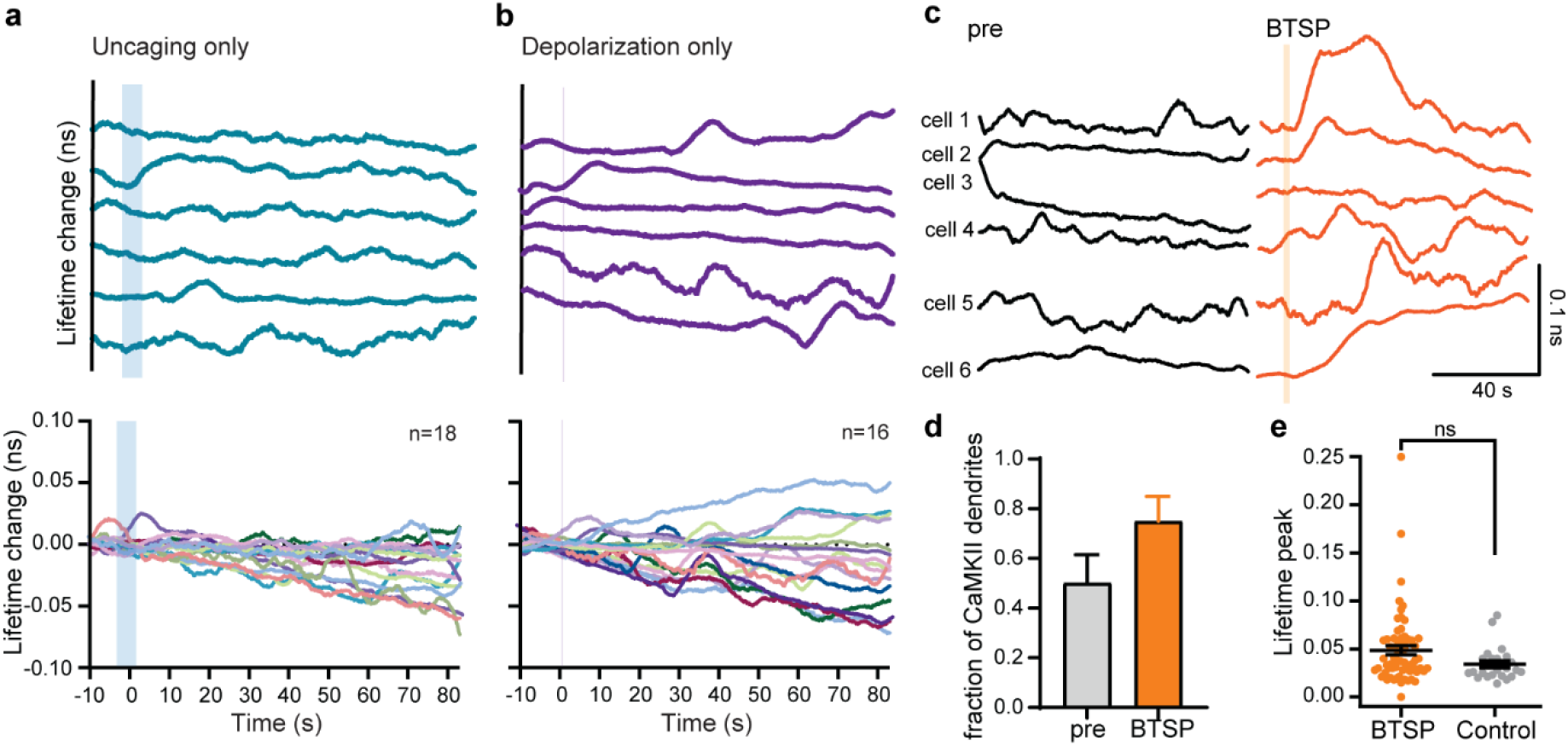
Control experiments show that delayed dendritic CaMKII is specific to BTSP induction. **a-b,** Top, Representative traces of CAMKII dendritic activity (smoothened by moving average of 60 points, 7.8 frames per seconds) of uncaging only and depolarization only experiments. Bottom, Lifetime change of all dendritic CaMKII activity for uncaging only (n=16) and depolarization only (n=19). The shaded region in representative and summary plots show where uncaging or depolarization was provided. **c,** Representative traces of smoothened Camuiα-2dv dendritic recording before (black) and after (orange) the BTSP protocol. **d,** Fraction of dendrites showing CaMKII activity in pre versus after induction of BTSP condition. **e,** Peak amplitude of CaMKII activation in responsive BTSP dendrites versus control dendrites (combined no-stim, uncaging only, and depolarization only) (p = 0.078, unpaired two-tailed t-test).

**Extended Figure 3-2:**
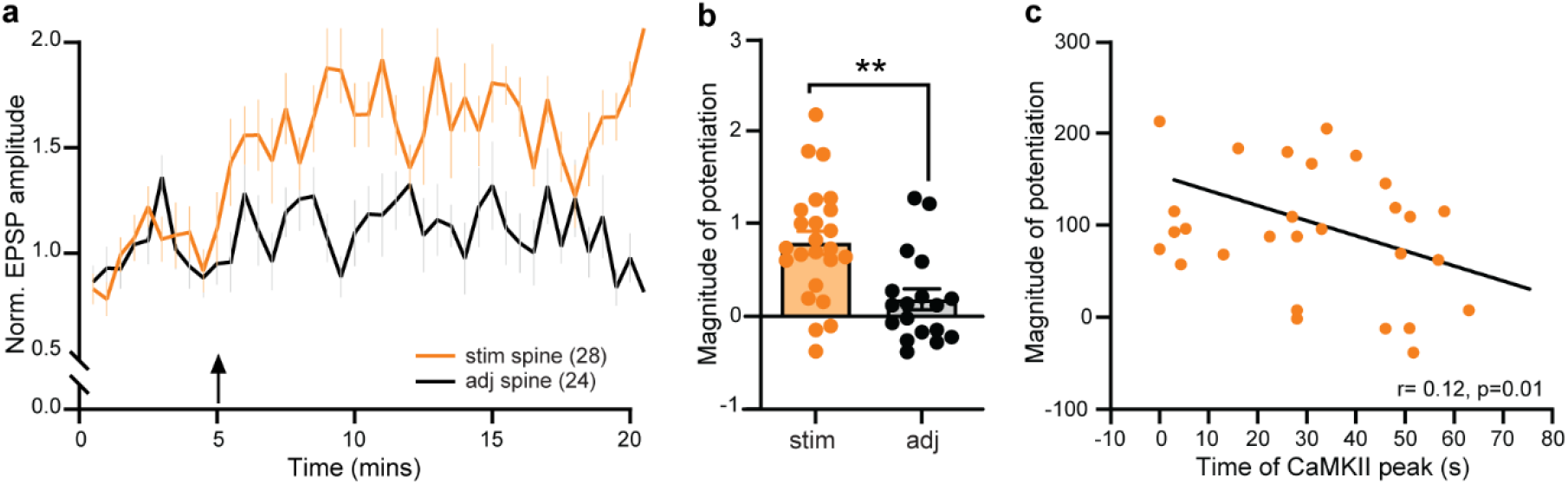
BTSP-induced synaptic potentiation is induced in 2dV-Camui labeled neurons. **a-b,** Averaged time course (**a**) and summary (25-30 min, **b**) of normalized EPSP amplitude in neurons expressing 2dV-Camui in the stimulated and adjacent spines. **p < 0.01, unpaired two-tailed t-test. **c,** Correlation between the magnitude of potentiation and the time of CaMKII peak after the BTSP protocol. The graph shows an inverse correlation such that earlier CaMKII activity results in a higher magnitude of potentiation (n=28 pairs, r^2^ = −0.12, p < 0.01).

**Extended Figure 3-3:**
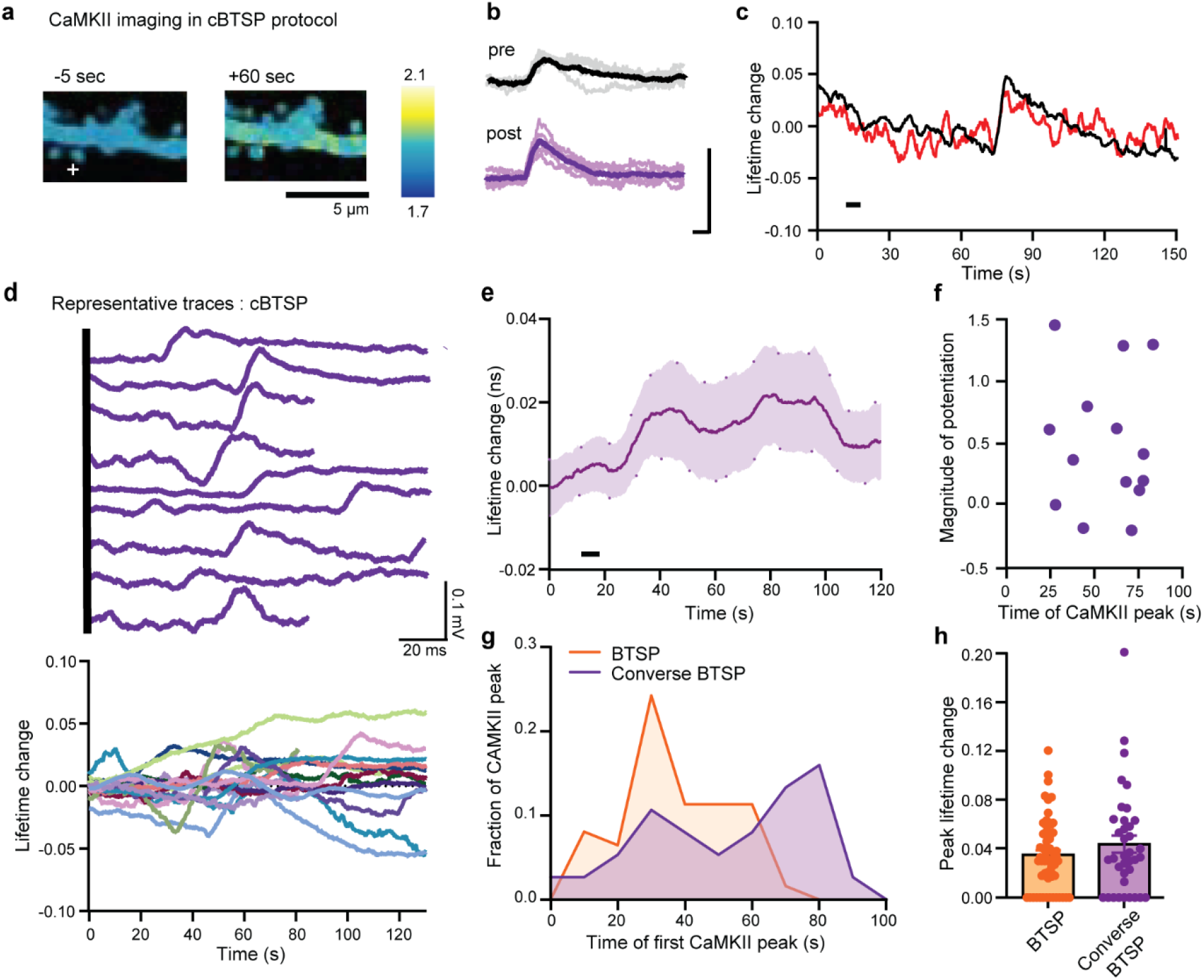
Converse BTSP protocol induces delayed dendritic and stochastic CaMKII (DDSC). **a,** Fluorescence lifetime images a 2dV-Camui expressing dendrite before, during and after the converse BTSP (cBTSP) protocol. **b,** Representative EPSP traces from stimulated spines before and after the cBTSP protocol. 10 traces and mean for each. **c,** Time courses of fluorescence lifetime changes in the stimulated spine and nearby dendrite from the dendrite in (**b**) (filtered). Black bar, cBTSP protocol. **d**, Top: Representative dendritic 2dV-Camui recordings. Bottom: All 2dV-Camui recordings (filtered) during uCBTSP induction. **e,** Mean + SEM lifetime change of dendritic 2dV-Camui recordings in response to cBTSP induction. **f,** Relationship between the magnitude of potentiation and the time of the first CaMKII peak after cBTSP protocol (n=15 pairs, r^2^=0.02, p=0.6). **g**, The frequency of CaMKII events (onset) as a function of time from the cBTSP protocol. **h,** Peak amplitude of CaMKII activation events before and after the cBTSP protocol.

**Extended Figure 3-4:**
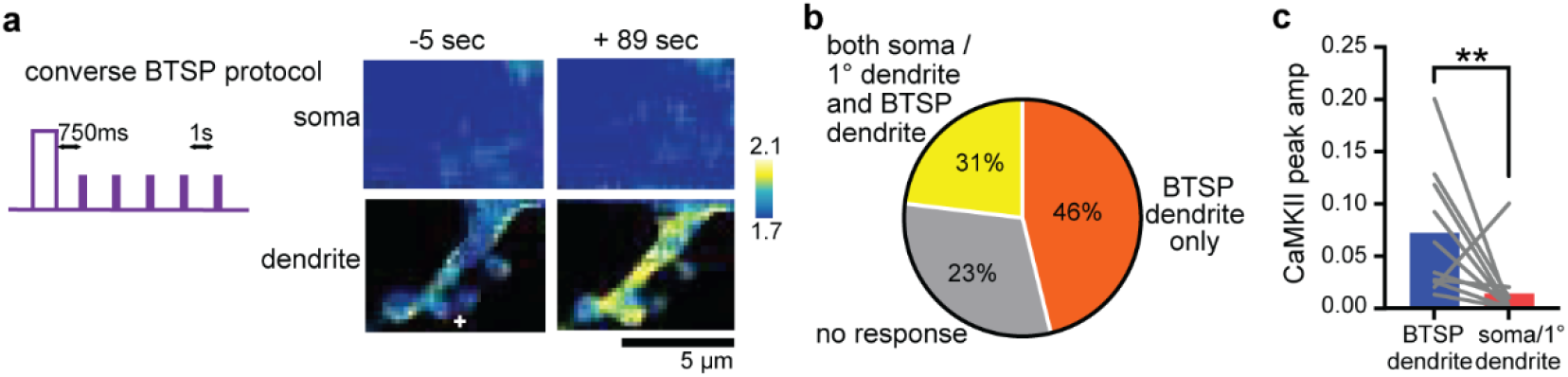
DDSC induced by converse BTSP protocol does not spread to the soma. **a,** Simultaneous FLIM imaging of 2dV-Camuiα in soma and stimulated dendrite during cBTSP. **b,** Pie chart for 14 recordings, out of which 46% showed an increase in CaMKII activity specifically in the dendrites but not in soma or primary dendrite. 31% of the recordings showed an increase in CaMKII activity in both the stimulated dendrite and the soma/primary dendrite and 23% of the dendrites showed no CaMKII activity. **c,** Peak amplitude of CaMKII activation events in the stimulated dendrite compared with that in the soma or the primary dendrite. **p<0.01. Paired two-tailed t-test.

**Extended Figure 4-1:**
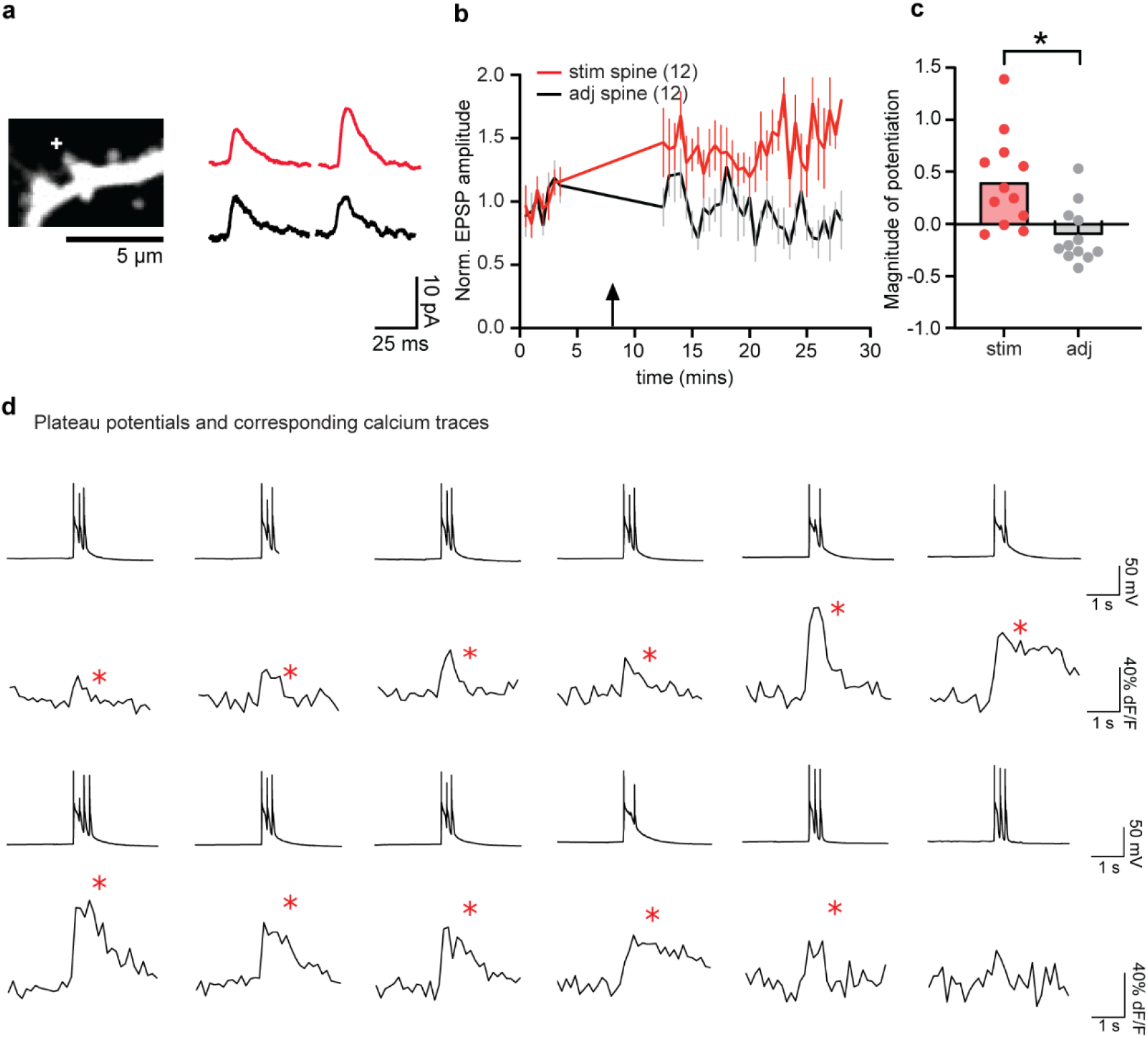
BTSP induces EPSP potentiation and plateau potentials observed during Ca^2+^ imaging experiments. **a,** A representative dendrite of a neuron loaded with Cal-590. Left: dendrite image, Right: EPSP traces before and after BTSP protocol. **b,** Averaged time course of normalized EPSP amplitude in stimulated and control spines. The gap in EPSP recording shows the time of calcium imaging (Fig. 4a**-c**). **c,** Group summary plot shows a higher magnitude of potentiation in stimulated spines (n=12) than adjacent spines (n=12, *p < 0.05, two-tailed t-test). **d**, Representative traces of voltage recordings showing that plateau potentials had corresponding Ca^2+^ traces (red *) in a majority of the examples (58/74, 3-standard deviation for Ca^2+^).

**Extended Figure 4-2:**
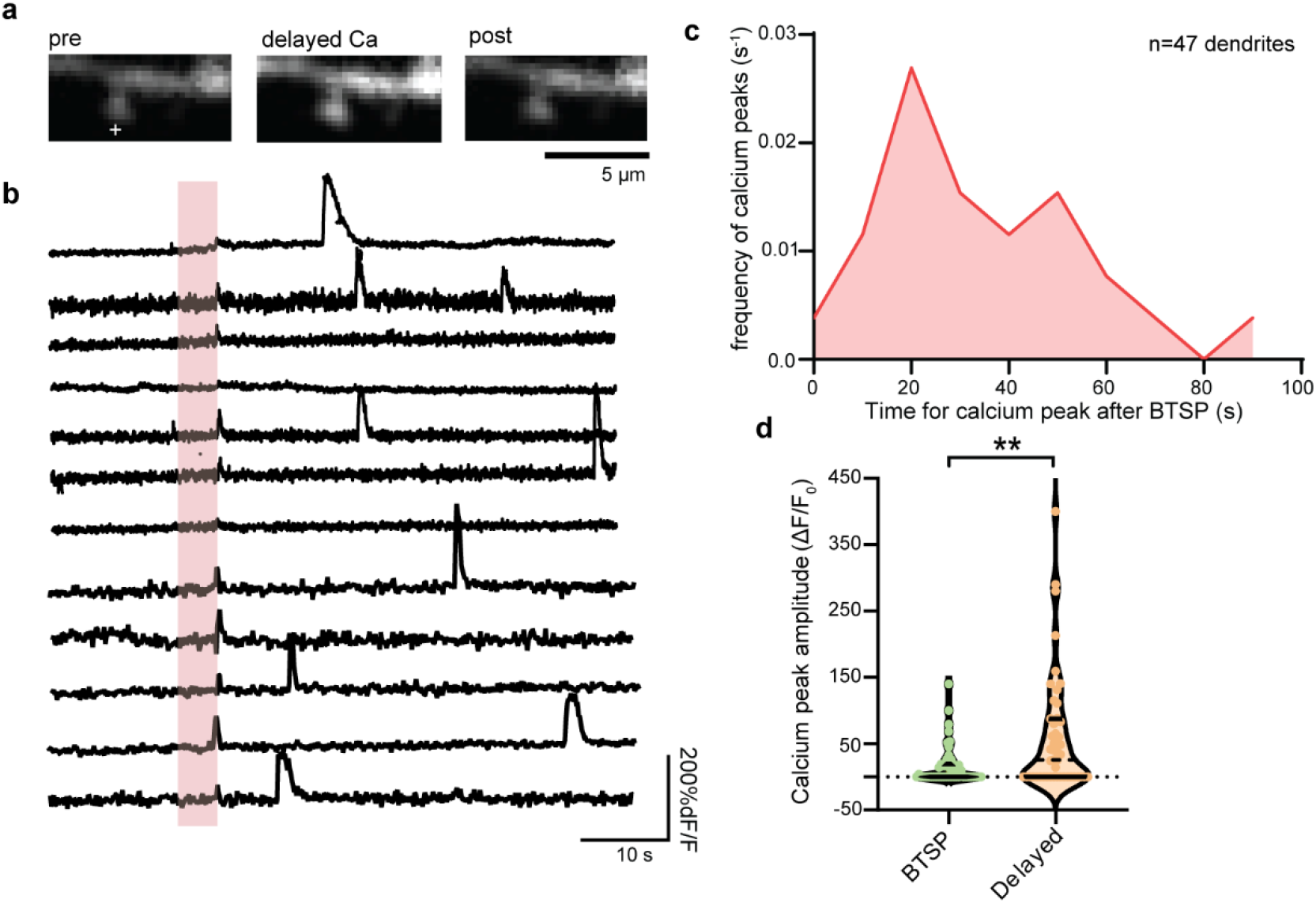
Characterization of delayed Ca^2+^ events in response to the BTSP protocol. **a,** Representative dendritic shaft filled with Cal 590 dye during a Ca^2+^ event. **b,** Representative dendritic calcium traces after the BTSP protocol (pink shadel. The traces show multiple calcium events after BTSP, in addition to smaller event during current injection of BTSP protocol. **c,** The frequency of Ca^2+^ events as a function of the time after the BTSP protocol. the frequency peak appears around ∼20-30 secs after the BTSP protocol. **d,** Ca^2+^ peak amplitude showed a smaller calcium during depolarization and a significantly larger delayed calcium peak amplitude, paired t-test, p<0.01.

## Notes

### Competing Interest Statement

The authors have declared no competing interest.

